# Unlocking *de novo* antibody design with generative artificial intelligence

**DOI:** 10.1101/2023.01.08.523187

**Authors:** Amir Shanehsazzadeh, Matt McPartlon, George Kasun, Andrea K. Steiger, John M. Sutton, Edriss Yassine, Cailen McCloskey, Robel Haile, Richard Shuai, Julian Alverio, Goran Rakocevic, Simon Levine, Jovan Cejovic, Jahir M. Gutierrez, Alex Morehead, Oleksii Dubrovskyi, Chelsea Chung, Breanna K. Luton, Nicolas Diaz, Christa Kohnert, Rebecca Consbruck, Hayley Carter, Chase LaCombe, Itti Bist, Phetsamay Vilaychack, Zahra Anderson, Lichen Xiu, Paul Bringas, Kimberly Alarcon, Bailey Knight, Macey Radach, Katherine Bateman, Gaelin Kopec-Belliveau, Dalton Chapman, Joshua Bennett, Abigail B. Ventura, Gustavo M. Canales, Muttappa Gowda, Kerianne A. Jackson, Rodante Caguiat, Amber Brown, Douglas Ganini da Silva, Zheyuan Guo, Shaheed Abdulhaqq, Lillian R. Klug, Miles Gander, Engin Yapici, Joshua Meier, Sharrol Bachas

## Abstract

Generative AI has the potential to redefine the process of therapeutic antibody discovery. In this report, we describe and validate deep generative models for the *de novo design* of antibodies against human epidermal growth factor receptor (HER2) without additional optimization. The models enabled an efficient workflow that combined *in silico* design methods with high-throughput experimental techniques to rapidly identify binders from a library of ∼10^6^ heavy chain complementarity-determining region (HCDR) variants. We demonstrated that the workflow achieves binding rates of 10.6% for HCDR3 and 1.8% for HCDR123 designs and is statistically superior to baselines. We further characterized 421 diverse binders using surface plasmon resonance (SPR), finding 71 with low nanomolar affinity similar to the therapeutic anti-HER2 antibody trastuzumab. A selected subset of 11 diverse high-affinity binders were functionally equivalent or superior to trastuzumab, with most demonstrating suitable developability features. We designed one binder with ∼3x higher cell-based potency compared to trastuzumab and another with improved cross-species reactivity^1^. Our generative AI approach unlocks an accelerated path to designing therapeutic antibodies against diverse targets.

## Introduction

Antibody drug development often begins with isolating initial leads by screening large libraries of random antibody variants against a target antigen. Techniques such as phage display^1^, yeast display^2^, immunization strategies, or single B-cell sequencing^3^ are typically employed to generate diverse binders. Further antibody optimization refines target affinity by focusing on the variable domains, which directly interact with antigens and are the key determinants of antibody affinity and specificity for targets^4–6^. Overall, these methods are laborious and often produce sub-optimal leads that fail during early-stage characterization. Computational approaches to *de novo* antibody design, that is, design against never-before-seen targets, have the potential to drastically reduce the time and resources necessary for therapeutic antibody development. The application of generative artificial intelligence (AI) methods for *de novo* antibody *design* is compelling, given the availability of large protein sequence and structure databases that can be leveraged for model training^7–19^.

Deep learning models have the potential to design antibodies with desirable properties^18, 20–25^. Several works have succeeded in optimizing antibodies using supervised learning approaches^26–29^. While different methods for AI-based de *novo* design of antibodies have been proposed with compelling *in silico* results, none have been coupled with high-throughput wet-lab validation^18, 20–25^. We exploited recent advances in DNA synthesis, recombinant antibody expression, fluorescence-activated cell sorting (FACS), high-throughput surface plasmon resonance (SPR), and next-generation sequencing (NGS) to experimentally screen and validate large libraries of antibody sequences. Combining these high-throughput techniques into optimized screening workflows of antibody designs obtained from generative AI could bridge the gap between computational antibody design and experimental validation.

Here, we developed generative AI models to produce antibody binders in a single round of design without further optimization cycles. Our models were trained on antibody-antigen complex structures and were constrained by removing known similar complexes to evaluate their capabilities for unbiased zero-shot *de novo design*. Zero-shot design, or deployment of models that have never seen a binder to a target antigen, is a relevant therapeutic problem based on its potential to unlock binder design against a broad set of novel targets without the need for further optimization. To assess the capability of our zero-shot *de novo* antibody design models, we used them to design heavy chain complementary determining regions (HCDRs) using trastuzumab and its target antigen HER2 as a model system^30, 31^. Following binder screening using our FACS-based Activity-specific Cell-Enrichment (ACE) Assay^TM^ and experimental validation using SPR, we identified diverse anti-HER2 binders with low nanomolar affinities, unique sequences, and distinct HCDR conformations. Finally, we reformatted a subset of designed binders as monoclonal IgG1 antibodies (mAbs) to assess potency and developability. In most cases, the high-affinity binders performed similarly to or, in some cases, better than trastuzumab. Taken together, this work paves the way toward fully *de novo* therapeutic antibody design using zero-shot generative AI to facilitate the development of biologics.

## Results

### Generative AI models for *de novo* antibody design built into an end-to-end screening and validation workflow

To assess the ability of our generative AI models to *de novo* design antibodies targeting specific antigens, we generated HCDR3 and HCDR123 sequences in a zero-shot fashion^6^. We followed the zero-shot definition as provided by language models GPT-3^32^ and ESM-1v^33^. We selected the therapeutic antibody trastuzumab, which targets HER2, as a template for the *de novo* design of HCDR sequences. The modeled HCDRs were conditioned on the HER2 antigen backbone structure derived from PDB:1N8Z (Chain C)^34^, trastuzumab’s framework sequences, and the trastuzumab-HER2 epitope. Neither the HCDRs nor the LCDRs of trastuzumab were provided as input to the models. Models were trained on the Structural Antibody Database (SAbDab)^11^. Any antibody in complex with the target antigen, HER2, or its homologs was removed from the training set. This was done by clustering the antibody-antigen complexes at 40% antigen sequence identity (Methods). We predicted the structure of the trastuzumab-HER2 complex using our structure design model, MaskedDesign. We then provided the predicted structure to our inverse folding model, IgMPNN, which designed HCDR sequences (Fig. 1, Methods). Sequence loss (model likelihood) was computed for the designed CDRs using IgMPNN and the predicted complex structure to obtain a score for ranking, according to which the top *k* sequences were sampled for subsequent wet-lab validation (Methods).

**Fig. 1.**
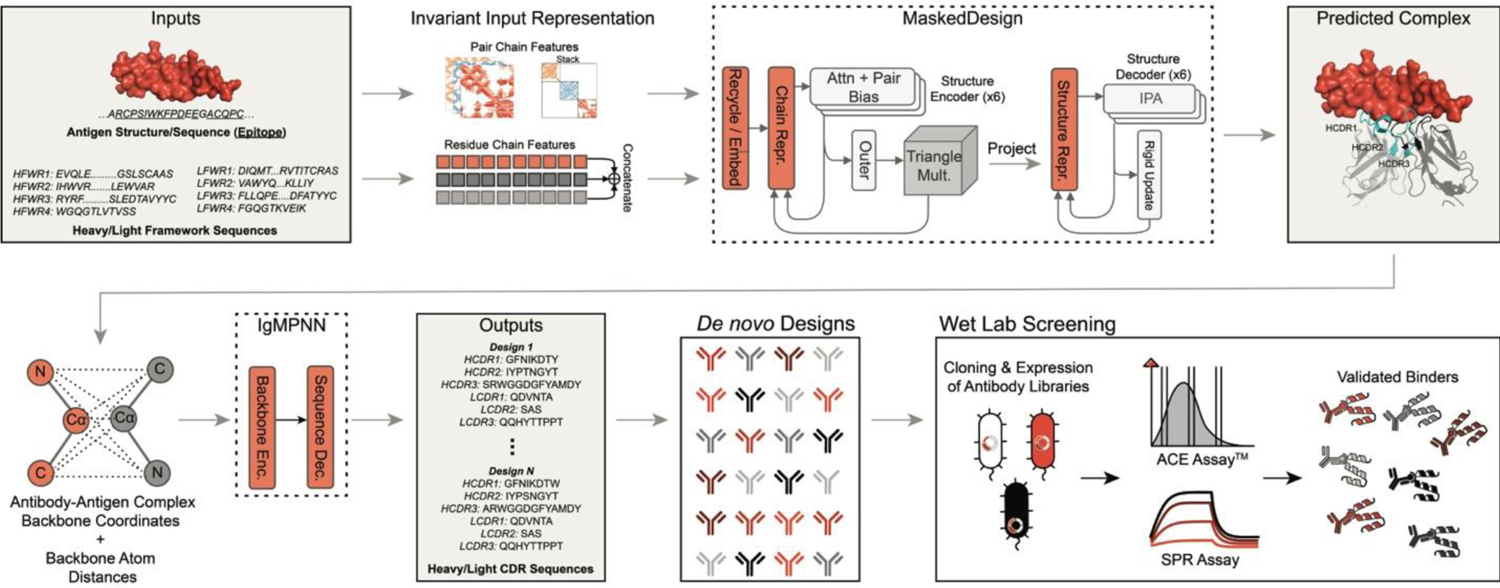
Zero-shot generative Al models for designing *de novo* antibodies. Deep learning models trained on antibody-antigen interactions combined with high-throughput wet-lab experimentation enabled the design of binders to an antigen never-before-seen by the models without further affinity maturation or lead optimization. Model architectures are depicted in dashed boxes. Model inputs and outputs are depicted with gray boxes in the background. Inputs to the model consisted of target antigen structure, target epitope region, and antibody framework sequences. None of the CDR sequences were provided to the models as input. Inputs are processed into invariant input representation and passed into the MaskedDesign model which predicts a docked antibody-antigen complex structure (sub-figure inspired by McPartlon et al.^16^). The predicted complex is passed to IgMPNN which designs CDRs (sub-figure inspired by Dauparas et al.^14^). *De novo* designed HCDRs are ordered as a library and are screened *in vitro* for binding.

Optimal HCDR lengths are unknown *a priori*, therefore, we sampled sequences with HCDR3 lengths of 9-17 residues, which represent 74.5% occurrence in the Observed Antibody Space (OAS)^10^. HCDR3 lengths were sampled according to the length distribution in OAS (Methods, Supplementary Table 1). We sampled HCDR1 and HCDR2 sequences consisting of 8 residues, as these are the most frequent lengths of HCDR1 (85.4% occurrence) and HCDR2 (64.8% occurrence) in OAS, and the lengths of trastuzumab’s HCDR1 and HCDR2 as defined by IMGT^35^.

We leveraged our FACS-based ACE Assay^TM^ (Methods) to screen fragment antigen-binding (Fab) libraries containing ∼10^6^ variants. In this assay, cloned antibody variants were evaluated for their antigen binding through intracellular expression in *E. coli*. The cells underwent permeabilization, staining for antibody expression and antigen binding, and isolation by FACS based on antigen-binding signal. NGS analysis was performed to compare the relative abundance of individual variants in the FACS-isolated populations to that of the original library. The significantly enriched variants were identified as the likely binders (Supplementary Fig. 1). To quantitatively validate the likely binders, we evaluated the performance of the ACE Assay^TM^ on a set of antibodies that were confirmed to be anti-HER2 binding or non-binding using SPR (Methods, Supplementary Fig. 2). Using this approach, we found that the ACE Assay^TM^ could classify SPR-measured positive and negative controls with nearly 95% precision and > 95% recall (Supplementary Fig. 1, Supplementary Tables 2-3). This robust workflow enabled the screening of a large variant population by the ACE Assay^TM^ with high precision (Fig. 1).

### Zero-shot *de novo* designs outperform biological baselines

We compared the binding rates of our generative AI models to relevant biological baselines derived by sampling sequences from OAS and SAbDab. We screened over 100,000 baseline sequences, including all HCDR3s and HCDR123s from SAbDab that matched appropriate length constraints, 50,000 HCDR3s and HCDR123s from OAS, and 10,000 HCDR123s from OAS which contained the same J gene as trastuzumab (Methods, Supplementary Table 4). Our models achieved a top 1,000 binding rate of 10.6% (HCDR3 design) and 1.8% (HCDR123 design), significantly outperforming the random OAS baseline by approximately 4-fold and 11-fold, respectively (Table 1). The models significantly outperformed the other baselines as well.

**Table 1.** Binding rates of AI *de novo* designs using correctly matched antigen (human HER2) as input vs. incorrect antigens (rat HER2, HER3, VEGF) as input and compared to biological baselines (OAS, OAS-J, SAbDab). For AI designs, the top 1,000 sequences by model likelihood are tested experimentally. AI designs with the correct antigen outperform all other strategies (indicated with * and bold; 2-sided Fisher’s exact *p* < 2 x 10^-4^, for all comparisons expect human HER2 vs. rat HER2 on HCDR123 with *p* = 0.0078).

| Method | Binding Rate (%) |  |
| --- | --- | --- |
|  | HCDR3 | HCDR123 |
| *De novo* (Human HER2) | **10.6\*** | **1.8\*** |
| Incorrect Antigen (Rat HER2) | 3.3 | 0.6 |
| Incorrect Antigen (Human HER3) | 2.4 | 0.2 |
| Incorrect Antigen (Human VEGF) | 2.5 | 0.0 |
| OAS | 2.77 | 0.17 |
| OAS-J | 5.41 | 0.32 |
| SAbDab | 3.26 | 0.07 |

The inverse folding model likelihoods were positively associated with binding rates, showing that sequences can be effectively prioritized in an unsupervised zero-shot manner. The designed sequences had increased binding rates when ranking by model likelihood, indicating that higher likelihood designs include more binders (Supplementary Tables 5-6). In fact, the top 100 binding rate was approximately 2-fold higher than the top 10,000 binding rate for both HCDR3 and HCDR123 design.

Furthermore, to show that the models meaningfully utilized antigen information, we designed off-target HCDRs conditioned on one of three incorrect antigens: rat HER2, human HER3, or human VEGF, instead of human HER2. We observed a significant performance decrease, with binding rates dropping greater than 3-fold, when designing sequences with incorrect antigens as input to the models (Table 1) and screening against human HER2.

### Generative models produce diverse binders

Analysis of the SPR-validated zero-shot AI designs (Fig. 2A) showed a binding affinity range to HER2 of over three orders of magnitude, with 71 exhibiting binding affinities less than 10 nM (Fig. 2B). Three binders showed tighter binding to HER2 than trastuzumab, with one binder displaying sub-nanomolar affinity. The HCDR3 length range was 11-15 residues (Supplementary Fig. 3A), a difference of ± 2 from trastuzumab. The HCDR3 sequences were also diverse from trastuzumab with edit distances of 2-12 (Fig. 2C). Interestingly, we detected binders that exhibited affinity less than 10 nM across all edit distances, suggesting paratopes similar to trastuzumab’s were not required for high affinity. Additionally, we observed higher diversity in HCDR3 centers, which correspond to the D gene, compared to the more conserved flanking V and J germline genes, indicating our models learned principles seen in natural immune repertoires (Fig. 2C)^36^. The binders were also dissimilar from one another, with a mean pairwise edit distance of 7.7 (Fig. 2D, Supplementary Fig. 3B). Despite the high sequence diversity, one potential explanation of the models’ success is the simple reproduction of training examples. The binders were distinct from any HCDRs in the training set and SAbDab (Fig. 2D, Supplementary Fig. 3C). Furthermore, when the binders were compared against OAS, an exponentially larger antibody database, some binders retained similarity to natural HCDRs (Fig. 2E) while most were distinct. These results indicate that the models were able to generate biologically relevant yet diverse HCDRs.

**Fig. 2.**
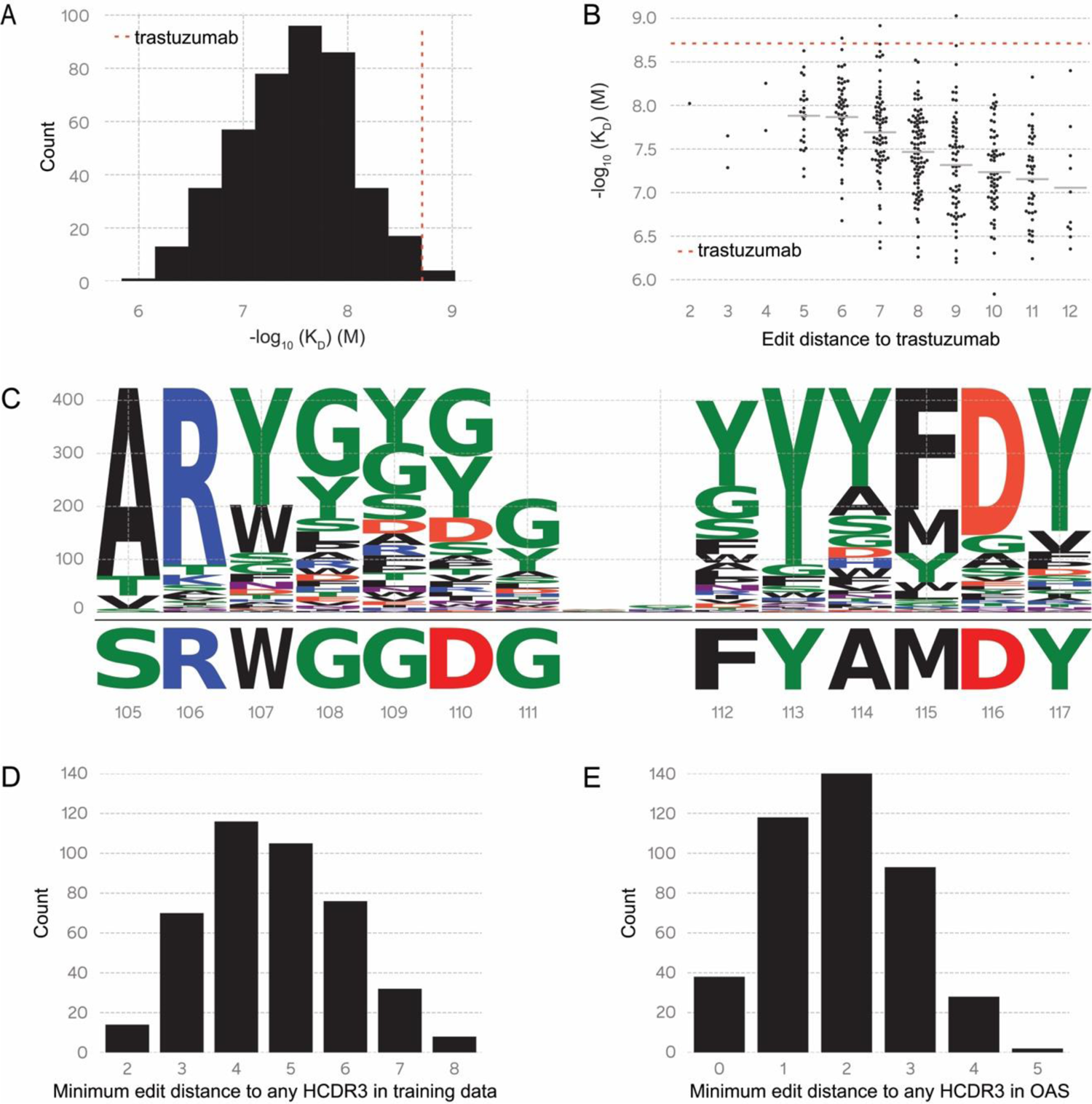
The design of diverse binders using zero-shot generative Al, further validated with SPR. **(A)** Plot of binder count vs SPR-measured binding affinity of *de novo* designed anti-HER2 binders (Fab format). The red dotted line represents the binding affinity (*-log_10_(K_D_) (M))* of trastuzumab. **(B)** SPR-measured binding affinities of *de novo* designed binders. The red dotted line represents the binding affinity of trastuzumab. Black dashes represent the mean for each edit distance bin. Edit distances range from 2 mutations (84.6% sequence identity) to 12 mutations (7.7% sequence identity). Each dot represents the mean affinity (*-log_10_(K_D_) (M))* across replicates. **(C)** Logo plot of HCDR3s of 421 SPR-validated binders compared to trastuzumab HCDR3 at the bottom. Greater diversity is observed in the centers of the designed HCDR3s. **(D)** Minimum edit distance of binders to training data HCDR3s (minimum of 2, maximum of 8, median of 5, mean of 4.68 ± 1.34 SD). **(E)** Minimum edit distance of binders to HCDR3s found in OAS (minimum of 0, maximum of 5, median of 2, mean of 1.91 ± 1.08 SD); 9.3% (38 out of 421) of the HCDR3 designs are found in OAS.

### Binders adopt variable binding mechanisms

We predicted structures for a diverse subset of the HCDR3 variants to better understand the structural basis of HER2 recognition (Methods). To this end, we built structural models of eight HER2-bound HCDR3 variants using the trastuzumab-HER2 Fab complex (PDB:1N8Z)^34^ as a starting template (Table 2, Supplementary Fig. 4). We performed locally constrained backbone geometry and side chain rotamer optimization followed by complex relaxation to correct global conformational ambiguities, steric clashes, and sub-optimal loop geometry^37^.

**Table 2.**
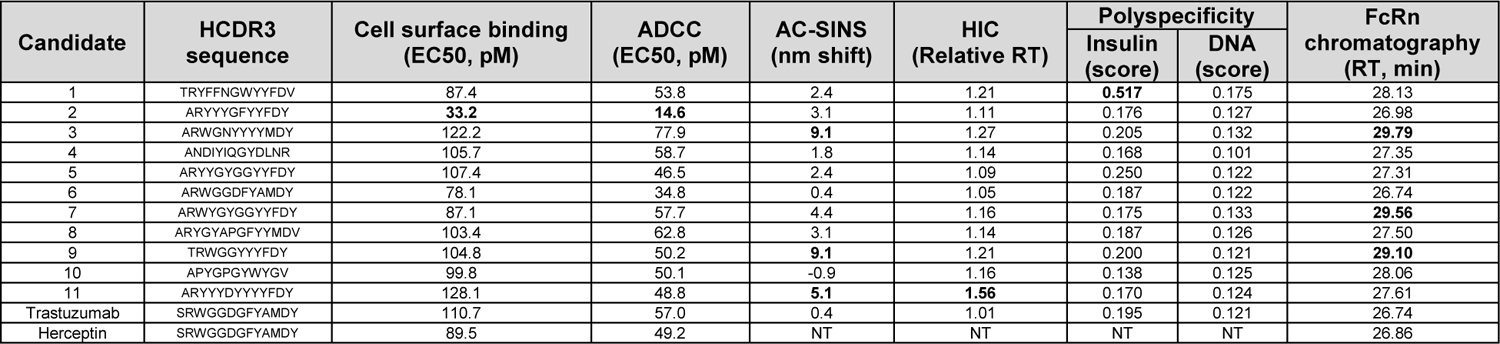
Binding and functional determination in cellular contents, developability and predictive pharmacokinetics of antibody candidates. Results for all 11 AI-designed antibody candidates for functional assessment by cell surface binding and ADCC activity in cell-based assays, developability by AC-SINS, HIC and polyspecificity, and predictive pharmacokinetics by FcRn chromatography. NT = not tested. Bolded values indicate those that exceed developability cutoffs for each assay.

Alignment of the designed HCDR3 regions with the trastuzumab-HER2 complex revealed a dynamic ensemble of conformations (Fig. 3, Supplementary Fig. 5). Antibody-aligned HCDR3 loop structural differences were broad, with local RMSD ranges of 1.1-6.8 Å when aligned across all HCDR3 atoms (Table 2). Despite sequence diversity, the eight *de novo* structural models were globally similar to trastuzumab with an all-atom HCDR3 RMSD range of 1.9-2.4 Å (Supplementary Fig. 5). In some variants, residues close to the HCDR3 sequence showed slight rotamer differences to account for longer loops or steric clashes from residues with larger side chains (Supplementary Fig. 6-7). Close analysis of the spatial orientation of the side chain conformations revealed identical side chains at five discrete spatial locations (Fig. 3). Two of these locations correspond to IMGT residue positions R106 and Y117 of trastuzumab, which are highly conserved in many antibodies^38^. In contrast, there was physiochemical conservation in positions W107, G109, and Y113 of trastuzumab, contributing to its paratope^30^. Although conserved spatially, these side chains originated from different positions, highlighting that conformational flexibility may be required for orienting key paratope residues.

**Fig. 3.**
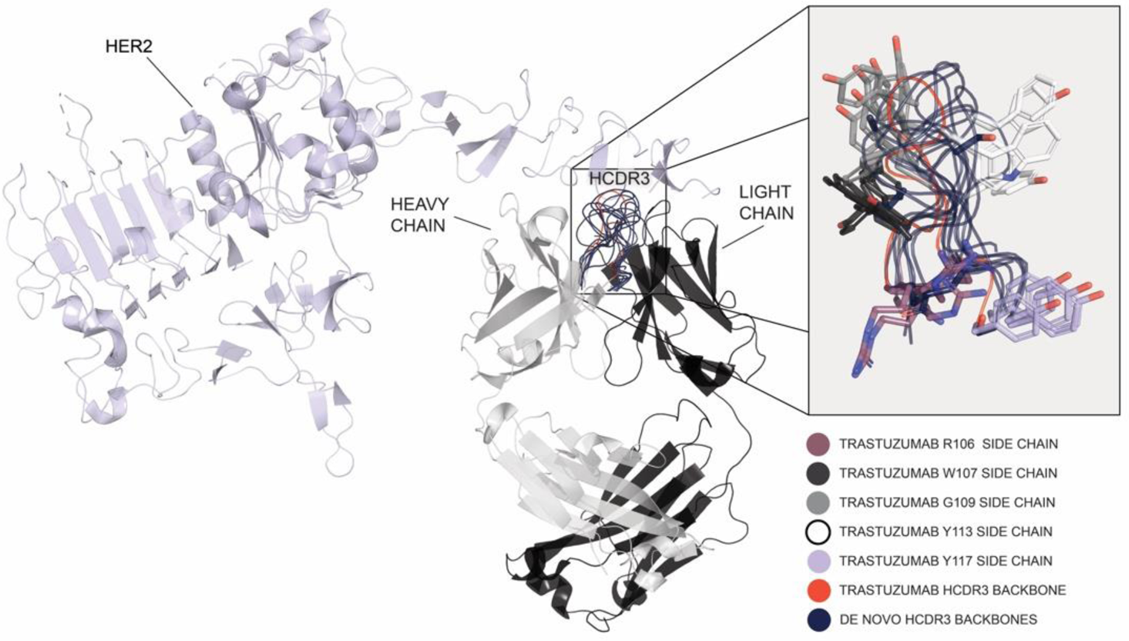
Structural model alignment of the designed HCDR3 regions with the trastuzumab-HER2 complex. Comparison of the trastuzumab-HER2 structure to the *de novo* designed binder complexes with HER2. Superimposition of the trastuzumab-HER2 structure (PDB:1N8Z) with *de novo* designed binder-HER2 complexes shows conformational differences in the HCDR3 backbone. Main chain backbone traces are depicted as ribbons and spatially conserved side chains are depicted as sticks. Despite the sequence and length diversity, key residues corresponding to the trastuzumab residues W107, G109, and Y113 are conserved in space (IMGT numbering scheme). Residues R106 and Y117 are also conserved although this is expected because of high conservation across most HCDR3s in immune repertoires.

Despite binding to the identical epitope, each variant exhibited distinct binding modes. Novel interactions not observed in the trastuzumab-HER2 complex were mostly formed between the designed HCDR3s and two HER2 domain IV surfaces (Supplementary Fig. 6-7). To further decipher binding determinants, we calculated the surface area buried by each HCDR3 variant when bound to HER2 (denoted as Interface, Supplementary Table 7). Several variants showed larger binding interface areas than trastuzumab. Interestingly, no significant correlation was observed between binding interface area and binding affinity. In addition, we calculated the grand average of hydropathy values (GRAVY)^39^ of each HCDR3 variant, a metric of their collective hydrophobic properties. We observed no significant correlation between affinity and hydropathy (Supplementary Table 7). Combined, these results suggest that the binding affinities of the designed HCDR3s were intrinsic to the sequences and were not driven by common physicochemical mechanisms, such as a potential bias towards increased hydrophobicity (Supplementary Table 7).

### *De novo* designed anti-HER2 binders display similar functional properties as trastuzumab

We explored the functionality and developability of 11 high affinity *de novo* designs. These variants were selected based on sequence diversity and biophysical properties and constrained to an affinity cutoff of < 10 nM. Trastuzumab served as a control. The variants were expressed as full-length monoclonal antibodies (mAbs) in mammalian CHO-K1 cells. The titers were equivalent to the trastuzumab titer; therefore, no sequence liabilities affected cellular expression. The mAbs were purified and analytical assays were conducted to assess homogeneity, purity, and concentration. Of the 11 variants, only one failed our > 97% purity cutoff by size exclusion chromatography (Supplementary Table 8). Intact mass by mass spectrometry showed uniform post-translational modifications in all variants. Biophysical characterization indicated all mAbs were monodispersed and had melting temperatures below trastuzumab but above the therapeutic cutoff of 60°C^40^ (Supplementary Table 8). Furthermore, mAb and Fab binding affinities were correlated (Supplementary Table 9), indicating a successful conversion into full-length mAbs, a limitation often observed with traditional techniques^41^.

Next, we probed the developability properties of each mAb using an array of assays that analyze hydrophobicity, self-aggregation, and polyspecificity (Methods, Table 2, Supplementary Fig. 8-11). Of the 11 mAbs, 7 displayed performances above the quality thresholds^40^. Only two mAbs displayed systematic failure across two or more assays. Neonatal Fc receptor (FcRn) chromatography is used as an initial *in vitro* method for evaluating the pharmacokinetic (PK) properties of antibodies. A reverse correlation between FcRn retention time and antibody half-life is typically expected^42^. Four of the 11 mAbs displayed subpar FcRn retention times, which could be attributed to poor biophysical properties such as self-aggregation (Table 2). These findings were expected since antibody variable domains have been previously shown to influence FcRn binding by IgGs^43^.

The mAb specificities were analyzed in a cross-reactivity SPR assay with HER2 paralogs and orthologs as antigens (Supplementary Fig. 12A). The *de novo* designed mAbs displayed similar species cross-reactivity profiles to trastuzumab, binding to human, cynomolgus, and canine HER2, with Candidate 2 outperforming all others, including trastuzumab (Supplementary Table 10). Non-specific binding was neither observed to mouse HER2 nor to the human paralogs HER1, HER3, and HER4 (Supplementary Fig. 12A, Supplementary Table 10). The candidate mAbs were tested for cell-surface antigen binding and antibody-dependent cell cytotoxicity activity using the HER2^+^ SK-OV-3 ovarian cancer cells (Supplementary Fig. 12B). We observed no significant binding differences between the *de novo*-designed mAbs and trastuzumab, except for Candidate 2 which displayed a ∼3-fold higher EC50 in the HER2^+^ cell surface binding assay. Compared to trastuzumab, similar or better cross-reactivity and functional profiles of the *de novo* designs indicate our approach enables diversification of mAbs optimized for developability and biological function (Table 2).

### Distinct interactions between HCDRs and epitope control functional properties

Improved cell-surface binding and cross-reactivity profiles could be linked to unique epitope interactions of the *de novo* HCDR3 designs. We utilized a high-throughput site-directed mutagenesis approach to map the binding contributions of each epitope residue to each mAb. We used a domain IV HER2 fragment as an epitope template and created alanine substitutions at residues within 5 Å of the paratope (Fig. 4A). The mutants were screened for mAb binding using an SPR assay, followed by ranking based on whether they were critical, partially critical, or not critical for binding (Methods). Construction of discrete epitope maps revealed that the mAb variants formed a network of unique interactions unrelated to trastuzumab, except for one epitope map (Supplementary Table 11). Interestingly, some HCDR3 variants formed critical interactions dominated by the light chain CDRs (Supplementary Table 11), implying significant conformational changes in the paratope and epitope facilitated novel interactions. Surface representations of epitope maps further highlighted similarities and differences in epitope interactions by each mAb, but also exemplified discrete epitope hotspots that influence function (Supplementary Fig. 13). The function of Candidates 2 and 3, which showed superior cell surface-binding potency and cross-reactivity, respectively, could be mapped to distinct hotspots based on differences to trastuzumab. Candidate 2 had a contribution switch at adjacent epitope residues K591 and D592, likely forming a hotspot for increased potency (Fig. 4B). Cynomolgus cross-reactivity was linked to distal regions of the HER2 domain IV epitope that mostly interact with the trastuzumab light chain (Fig. 4A). Candidate 3 led to considerable contribution changes in epitope residues F595, M611, and I613, likely forming a hotspot for cross-reactivity, because they are structurally close to the two polymorphic differences between cynomolgus and human HER2 (Fig. 4B, Supplementary Table 11). Together, these results highlight the versatile, functional properties that can be obtained from diverse HCDR3 designs (Fig. 4C).

**Fig. 4.**
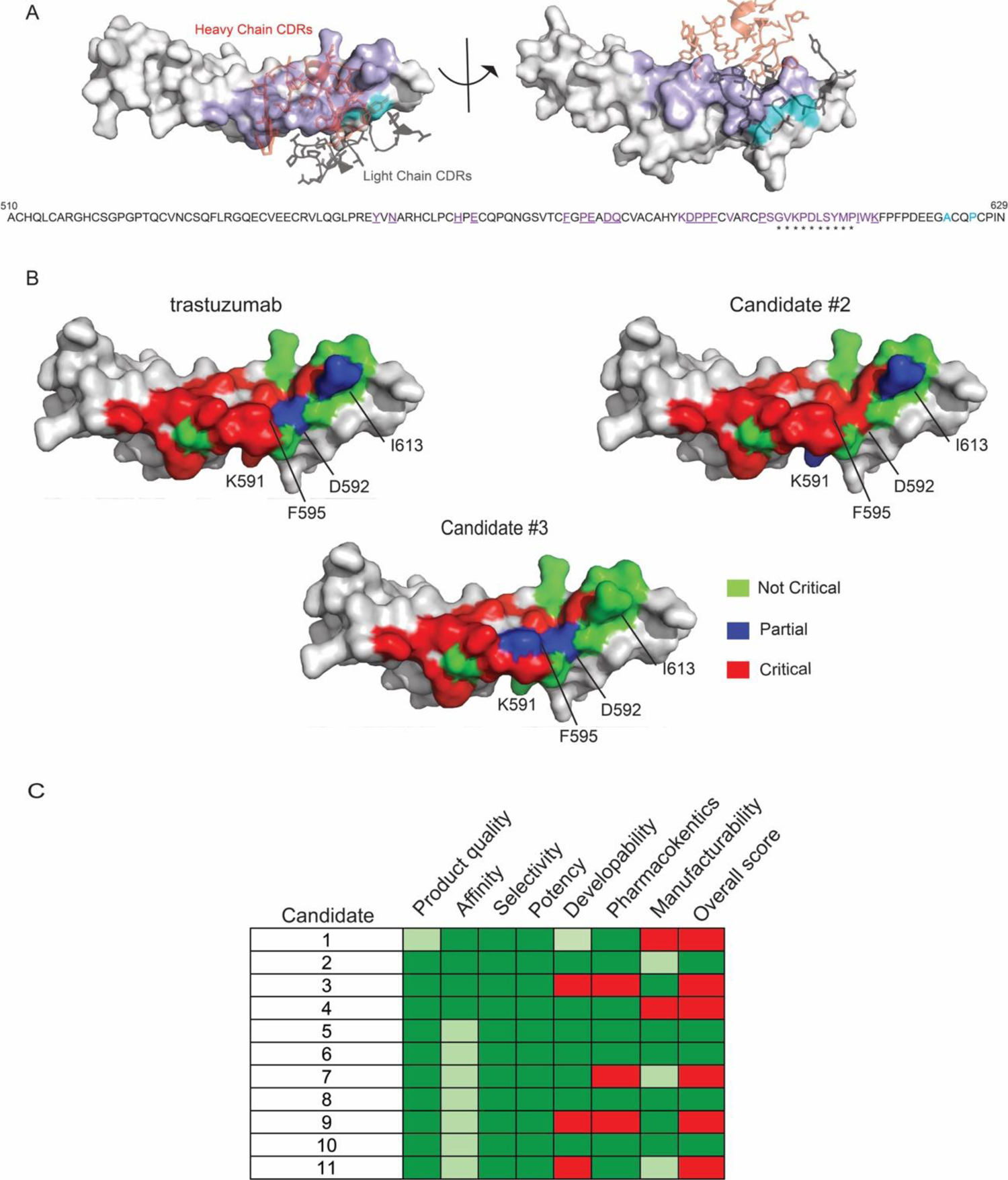
Functional properties of 11 candidate mAbs. **(A)** Residues that constitute the HER2 Domain IV epitope fragment used in alanine scanning mutagenesis experiments. Residues within 5 A of trastuzumab heavy and light chain are colored purple. Epitope residues that contact the heavy chain are underlined. Residue positions that differ between human and cynomolgus epitope are colored cyan. Heavy chain and light chain CDRs are colored red and gray, respectively. **(B)** Surface representation of the epitope maps of trastuzumab, high potency candidate 2, and most cross-reactive candidate 3. Residue surfaces are colored according to their contributions to binding in the figure legend. Hotspot residues that contribute to function are highlighted. **(C)** Summary of functional properties of designed mAbs.

## Discussion

A particularly difficult aspect of antibody drug design is the initial step of lead candidate identification due to the labor-intensive and uncontrolled nature of traditional discovery and screening methods^1–6^. Computational antibody design strategies offer alternative solutions^18, 20–23^. Of particular interest is epitope-specific binder design that can be achieved under controllable *in silico* settings and be subsequently validated experimentally. Here, we demonstrated the potential of zero-shot generative AI models to reduce the need for repeated library screening or animal immunization, which yield leads that typically need optimization and are not necessarily epitope-specific. Our study yielded novel antibodies that bound human HER2 with other characteristics comparable to and, in some cases, superior to the therapeutic antibody trastuzumab. In addition, our models generated *de novo* HCDR3 and HCDR123 designs with significantly greater efficiency than relevant biological baselines, enabling a larger and more diverse pool of leads for candidate selection. Importantly, the AI-generated sequences were distinct from sequences in the training set and the vast majority of OAS suggesting the models are not simply memorizing the training data. Structural modeling of a subset of binders revealed high backbone conformational variability, but preservation of important positional interactions with the HER2 antigen. Altogether, the observed high diversity in the sequences and conformations of the antigen-binding variable regions indicates that our models recapitulate natural immunity principles while generalizing to new sequences.

Cell expression of high-affinity anti-HER2 antibodies incorporating *de novo* HCDR3 designs resulted in functional and developability properties similar to or better than trastuzumab. These were linked to HER2 epitope hotspots that control binding potency and cross-reactivity. We found that our designed antibodies form unique interactions with epitope residues distinct from the overall interaction map of trastuzumab. In conclusion, the performance of our models indicates a propensity to design CDRs with favorable antibody-like biophysical properties and targeting of the selected epitope, enabling the mitigation of downstream developability risks.

Building on the demonstrated progress, future work could extend generative AI models to enable the *de novo* design of all CDRs and framework regions against more antigens and epitopes. If generalizable, the epitope-specific targeting of antigens of interest during antibody design can improve efficacy or enable novel pharmacology. In addition to further diversifying possible binding solutions, novel frameworks may result in antibodies with better clinical-grade properties, including developability and functionality. In conclusion, generative AI models combined with high-throughput wet-lab screening capabilities and followed by the selection of final drug candidates has the potential of unlocking new capabilities in the field of therapeutic antibody design.

## Acknowledgements

The authors wish to thank Matthew Weinstock, David A. Spencer, Luka Stojanovic, and Alec Jaeger for early discussions; Christian Stegmann, Jens Plassmeier, Christine Lemke, Mario Sanches, Zachary McDargh, Bradley Emi, Thomas Wrona, Sarah Korman, Byron Olsen, Zach Jonasson, Joseph Sirosh, Ivana Magovcevic-Liebisch, Dan Rabinovitsj, Daniele Biasci, Joerg Stelling, Victor Greiff, and Amaro Taylor-Weiner for critical review of this manuscript; Joe Kaiser for providing MLOps and DevOps support; Stephanie Yasko and Marcin Klapczynski for schematics support; Katherine Moran for program management support; Greg Schiffman, Andreas Busch, and Sean McClain for continual support; and Kfir Lapid for editorial assistance.

## Author Contributions

A.S. and S.B. wrote the manuscript and prepared figures with contributions from all authors. A.S., M.M., and J.M. devised strategies for implementing generative AI models for antibody design. A.S., M.M., R.S., J.A., G.R., S.L., J.C., J.M.G., A.M., J.M. supported AI efforts for antibody design. A.K.S., J.M.S., C.K., Re.C., H.C., A.B.V., J.B., K.A.J., S.A., and Mi.G. designed and conducted flow cytometry and NGS experiments related to the ACE Assay^TM^. C.C., B.K.L, N.D., A.B., G.K., Ed.Y., and S.B. designed and conducted SPR validation experiments. C.L., I.B., P.V., Z.A., L.X., P.B., D.G.S., designed and conducted developability experiments and generated quality assessment data on candidate mAbs. O.D., K.A., and En.Y. designed and conducted cell-based functionality experiments on candidate mAbs. K.B., G.K.B, D.C., and Z.G. designed vectors and conducted cloning of libraries and variants for ACE Assay^TM^ and SPR studies. B.K., M.R., G.M.C., Mu.G., and Ro.C. designed and conducted antibody expression and cultivation experiments. G.K., Ed.Y., C.M., R.H., and S.B designed and conducted epitope mapping, cross-reactivity, and biophysical characterization experiments. Ed.Y. and S.B. designed IgG1 mAbs for developability experiments. Structural analysis and calculations were performed by S.B. with support from A.S. and M.M. L.R.K, S.A., Mi.G., En.Y., J.M., A.S., and S.B. were responsible for project governance.

## Methods

### Antibody Numbering and CDR Definitions

IMGT numbering and CDR definitions are used throughout this work^1^.

### *In Silico* Binder Design

We designed sequences *in silico* through a two-step process. In the first step, MaskedDesign, our structure design model, was given the antigen sequence and structure, the position of the epitope residues on the antigen, and the antibody framework sequences. MaskedDesign predicted the 3D backbone structure of a bound antibody-antigen complex. This predicted complex structure was then given to IgMPNN, our inverse folding model. IgMPNN predicted HCDR sequences that will form the given complex, which are our designed sequences. For each designed sequence, we computed the sequence loss (model log-likelihood) using IgMPNN and the predicted backbone complex structure as a measure of binding likelihood. Finally, we ranked sequences using these scores and selected the top k sequences for *in vitro* validation.

### Structure Design Model (MaskedDesign)

MaskedDesign is similar to the model described in McPartlon et al^2^.

#### Model Inputs

MaskedDesign took an annotated structure as input. During training, we provided:

- The sequence and the backbone coordinates of the antigen.
- A binary mask indicating the epitope residues.
- The sequence of the heavy chain of the antibody (backbone atom coordinates and the sequences of the CDRs were designed and were masked out).
- The sequence of the light chain of the antibody, if a light chain exists (backbone atom coordinates and the sequences of the CDRs were designed and were masked out).

From this input, we computed residue and pairwise features.

#### Residue Features

- **Residue type:** One-hot sequence encoding for the antigen and antibody chains with 20 tokens for natural amino acids and a <mask> token for masked sub-sequences and missing values.
- **Chain encoding:** We enumerated the protein chains in the structure and one-hot encoded the values for each residue.
- **Epitope:** Binary (0/1) tensor indicating the sampled epitope residues.
- **Dihedral angle encoding:** We calculated the dihedral angles along the backbone of each chain (that did not have its coordinates masked). We then binned the angles into 18 equidistant 20° bins, along with bins for masked values or missing values, and calculated a one-hot encoding of these bins. If the coordinates of residue i are masked, then the angles cp_i-1_ and 1/_i+1_ are masked.

### Pairwise Features

- **Relative sequence separation:** We calculated the signed distance between each pair of residues within a chain (the number of residues between them, positive if the first residue is downstream of the second one). We then binned the values into 29 bins: 14 positive bins (0, 1, 2, 3, 4, 5, 6, 7, 8, 10, 10-12, 12-15, 15-20, 20-30, and > 30); 14 negative bins matching the positive bins; and a bin for the value 0, along with a bin for masked values. We then calculated a one-hot encoding of these bins.
- **Pairwise distance encoding:** For each pair of residues whose coordinates were not masked, we calculated the distances between the Ca and all four backbone atoms; we then binned the calculated distances into seven bins (< 2 Å, 2-4 Å, 4-6 Å, 6-8 Å, 8-10 Å, 10-12 Å, and > 12 Å). Finally, we calculated a one-hot encoding of these bins.
- **Pairwise interaction flag encoding:** Binary (0/1) tensor indicating if the two residues were in contact (Ca atoms are within 10 Å).

<u>Note:</u> This is an optional feature that is not used for the design model (neither in training nor inference) but is used for the denoising model.
- **trRosetta orientation:** We calculated the 0, w, 1/ orientation as described in Du et al^3^. We then binned the values into 18 equidistant 20° bins, along with bins for masked values or missing values, and one-hot encoded the values.

We concatenated these features to form the residue and pair input tensors.

### Model Architecture

Our structure design model consisted of two major parts, which we labeled as the encoder and the decoder.

Each encoder layer was made up of two blocks: a residue update block (which updates the residue embeddings), and a residue pair update block (which updates the pairwise “interaction” embeddings):

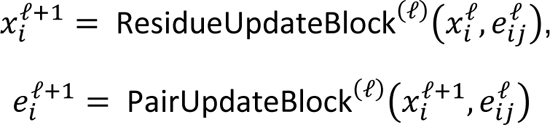

A residue update block performed a multi-head attention update with pair bias and residual connections, followed by dropout, layer normalization, and a fully connected layer that also has residual connections. A pair update block performed incoming and outgoing triangle-multiplicative updates.

The decoder layers only performed residue embedding updates. The architecture of the decoder layer residue update block was the same as that of the encoder residue update block, with the difference of shared weights across all layers of the decoder. We passed the residue embeddings from the decoder layer through a linear projection layer to make the intermediate (from states of the intermediate decoder layers) and final (from the state of the final decoder layer) coordinate predictions.

The output of the final decoder layer was also passed through an additional fully connected layer, the outputs of which were used as logits for an auxiliary sequence prediction loss.

### Loss Function

We applied the following loss function during the training of the structure design model:

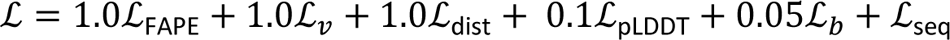

L_FAPE_ is the FAPE loss, as defined by Jumper et al^4^. We computed FAPE over the final output from the model and all intermediate decoder layers.

L_v_ is a steric violation loss, which was based on Ca-Ca distances between each pair of residues and is defined as: L_v_(x) = max(0,2.7 - x).

L_dist_ is a binned pairwise distance loss. We used the pair features from the model and passed them through a fully connected layer to compute the prediction logits. The labels were calculated by binning the pairwise distances for four atom pairs (Ca, X) where X E {N, Ca, C, C/3} into bins of 0.4 Å width in the range from 2.5 Å to 20 Å, and one extra bin for distances greater than 20. We computed the loss value by comparing the predictions to the ground-truth labels and applying cross-entropy to the result.

L_pLDDT_ was calculated by passing the outputs of the final decoder layer through a fully connected layer with output representing 20 equal-width binned log-likelihoods in the range [0, 1]. We computed the loss value by comparing the predictions to the ground-truth labels and applying cross-entropy to the result.

L_b_ is the backbone bond length loss, which we computed as the Huber loss calculated over the distances between the consecutive backbone atoms along the backbone of each chain in the input structure.

L_seq_ is the auxiliary cross-entropy sequence loss, applied over the masked-out part of the sequence.

### Hyperparameters and Training

We trained the structure model with the following dimensions: All encoder layers and decoder blocks used hidden layers of size 196 for the residue embeddings and 64 for the pairwise embeddings. Both the encoder and the decoder contained 5 layers. During training, the dropout rate was set to 0.1.

We trained the model on antibody-antigen complexes in SAbDab with the HER2 cluster removed and antibodies with the same HCDR3 as trastuzumab removed (details on the curation of datasets are in the Antibody Databases section). We used the Adam optimizer, with a maximum learning rate of 0.001, linear warm-up for 500 steps, followed by linear learning rate decay 1 x 10^-5^ over the rest of the training steps.

### CDR Structure Design

At inference time, the model took as input the structure and sequence of the antigen, the selection of desired epitope residues, the sequences of the antibody framework regions, and the lengths of the CDR regions to be designed. The model outputs a 3D structure of the antibody-antigen complex with CDRs having specified lengths. In addition to a design model, we also trained a denoising model which took as input an antibody-antigen complex structure, added noise, and recovered or “denoised” the complex structure. The denoising model was trained with the same architecture, data, hyperparameters, and training scheme as the design model. The generated structure from the design model was passed into the denoising model and the denoised structure is then passed to the inverse folding model to design the antibody CDRs.

### Inverse Folding Model (IgMPNN)

IgMPNN is similar to what is described in Shanehsazzadeh et al^5^. IgMPNN uses a message-passing neural network (MPNN) scheme^6^. Our implementation took as input the 3D coordinates of backbone residues of an antibody-antigen complex. We defined a protein graph as a directed graph where residues were represented as nodes and shared edges with their *k* nearest neighbors. We used *k = 48* in all of our experiments. We initialized the node features *x_i_* and edge features *e^t=O^* in our protein graph using the following features^6^:

- Distances between *Ca*-*Ca* atoms
- Relative *Ca*-*Ca*-*Ca* frame orientations and rotations
- Backbone dihedral angles
- Binary (0/1) features that determine relative chain positions
- Relative position encodings

Our featurization differed from the original MPNN^6^ in that we did not assume access to any side chain atoms, and thus we did not include any pairwise distance features involving sidechain atoms.

Our initial features *x_O_* then get passed into our message passing neural network encoder. Our network had multiple message-passing phases, during which the hidden state of each node in the graph *h^t^* and edge *e^t^* was updated according to:

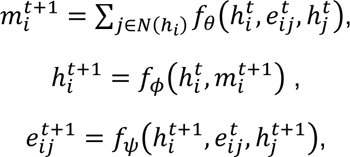

where *f_0_*is our message update function, *f_¢_* is our node update function, *f_l/J_* is our edge update function, and *N(h_i_)* is the neighboring nodes for a given node *i* in the graph. We used 128 as our hidden node and edge dimensionality size throughout the network. IgMPNN utilized three encoder layers, performing message passing, node updates, and edge updates. The output was then fed into the decoder, which had three layers. The decoder performed message passing to update the node representations according to:

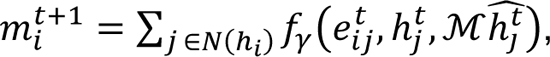

where *f_y_* is our decoder message update function. The ground truth context, *h^-t^*, which is an embedding of the ground truth residues, was provided as input only when it was allowed by our causal decoding mask *M*. The decoder is masked to prevent the model from incorporating information from nodes that have yet to be decoded while allowing the decoder to access information from nodes that have already been decoded. When decoding a given residue during training time, instead of accessing the embeddings for predicted residues at previously decoded positions, the decoder accessed embeddings for ground truth residues at previously decoded positions. The model decoded antibody CDRs in sequential order during training: HCDR1, HCDR2, HCDR3, LCDR1, LCDR2, LCDR3 (or only HCDR1, HCDR2, HCDR3 if no light chain is present). During inference, a different order of CDRs can be specified. Within each CDR, the decoding order is random. Finally, we projected our final node embeddings to logits.

We trained the model using cross-entropy loss. The model is first pretrained on proteins from the PDB curated into the “General PDB” dataset (details on the curation of datasets are in the Antibody Databases section). The model is then fine-tuned on antibody-antigen complexes in SAbDab with the HER2 cluster removed and antibodies with the same HCDR3 as trastuzumab removed (details on the curation of datasets are in the Antibody Databases section). We used the Adam optimizer, with a maximum learning rate of 0.001, linear warm-up for 500 steps, followed by linear learning rate decay 1e^-5^ over the rest of the training steps.

At inference time, the model is given an antibody-antigen complex structure, such as a predicted structure from MaskedDesign, and predicts the CDRs of the antibody. We predicted HCDR3 first, followed by HCDR1 and HCDR2 (and then the LCDRs, which are kept fixed during *in vitro* screening, in numerical order). We selected this order to prevent the model from conditioning on potentially false CDR predictions when predicting HCDR3. We note that the model is never provided ground truth CDR sequences at inference time. For sampling, we used weighted random sampling after applying softmax with a temperature of T = 0.5 to the model’s logits. We also prevented the model from sampling cysteines to avoid having unpaired cysteines in the CDRs. Predicted CDRs are then scored by passing the sequence back into the model and computing log-likelihood over the relevant tokens and positions. This score is used to rank the sequences and select the top ones for *in vitro* screening.

### Antibody Databases

The Observed Antibody Space (OAS)^7^ was retrieved on February 1^st^, 2022. The Structural Antibody Database (SAbDab)^8^ was retrieved on December 6^th^, 2022. The corresponding PDB files were downloaded from RCSB PDB^9^. To generate a high-quality dataset used for training, we applied the following filters:

- Drop entries without a heavy antibody chain.
- Drop entries without an antigen.
- Drop entries where the PDB id, heavy, light, and antigen chains were repeated (duplicated).
- Drop entries where one of the HCDRs was too short (shorter than 5 amino acids for HCDR1 and HCDR2, and shorter than 7 amino acids for HCDR3).
- Drop entries where one of the HCDRs was too long (longer than 26 amino acids for any of the HCDRs).
- Drop entries where more than 10% of heavy chain residues were missing from the structure.
- Drop entries where more than 25% of antigen residues were missing from the structure.

After filtering, there were 6933 entries left in the database. We then applied sequence clustering to the antigen sequences using mmseqs2^10^ version 13.45111, with the following parameters: min-seq-id=0.4, cov-mode=1, cluster-mode=2, cluster-reassign=true. We used these cluster annotations when splitting the data into train, validation, and test folds, assigning an entire cluster to one of the three subsets.

The General PDB dataset was created from a selection of PDB files available in the RSCB PDB^9^ database. We made the selection using the Graphein library (version 1.0)^11^, downloading all PDB structures where each chain was longer than 40 and shorter than 500 amino acids. We further filtered out any structures that contained chains with missing backbone atoms. This resulted in a dataset with 74734 entries. We then applied sequence clustering to the antigen sequences using mmseqs2 with the same set of parameters used for clustering the SAbDab data (min-seq-id=0.4, cov-mode=1, cluster-mode=2, cluster-reassign=true). As with SAbDab, we used these cluster annotations when splitting the data into train, validation, and test folds, assigning an entire cluster to one of the three subsets.

To compute edit distance (number of mutations) between antibody sequences, we used the Levenshtein distance, denoted Lev. The “Minimum HCDR3 edit distance to OAS” was computed by taking the minimum edit distance between an HCDR3 of interest and all HCDR3s in OAS:

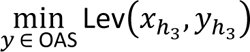

This value was analogously computed for HCDR1 and HCDR2 as well as for other databases such as SAbDab or our training data. The “Minimum HCDR123 edit distance to OAS” was computed by taking the minimum edit distance between the tuple of HCDRs (HCDR1, HCDR2, HCDR3) belonging to an antibody of interest and all such tuples in OAS, analogously computed for other databases:

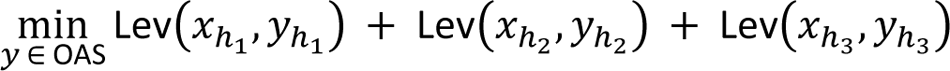

To compute the OAS HCDR3 length distribution (Supplementary Table 1), we iterated through all heavy chain sequences in OAS, considered the HCDR3 length, and maintained a tally of HCDR3 lengths. We then restricted to HCDR3 sequences with lengths between 9 and 17 amino acids, normalized to get the length distribution. For the OAS-J HCDR3 length distribution, we performed an analogous process but iterated only through heavy chain sequences in OAS that had trastuzumab’s J-gene. For the SAbDab HCDR3 length distribution, we took all unique HCDR3 sequences in SAbDab with lengths between 9 and 17 amino acids and computed the frequencies at each length.

### Model Structural Inputs

As input to the models, we provided an antigen structure and specified an epitope. For the *de novo* setting, we provided the structure of human HER2 from PDB:1N8Z (Chain C)^12^ and specified the trastuzumab epitope determined from the structure (i.e., antigen residues within 5 Å of the antibody with distance computed over all atoms). In this setting, we successfully designed using versions of the antigen structure with variable amounts of noise or having been relaxed with Rosetta^a^.

To show the model’s dependence on the antigen information, we attempted to design human HER2 binders with three incorrect antigens, namely rat HER2, HER3, and VEGF. For rat HER2, we used the structure from PDB:1N8Y (Chain A)^12^. For HER3, we used the structure from PDB:7MN8 (Chain A)^13^. For VEGF, we used the structure from PDB:1CZ8 (Chains A, B)^14^. For rat HER2 and HER3, we specified an epitope based on sequential/structural homology to the trastuzumab epitope of human HER2. For VEGF, we specified the ranibizumab^15^ epitope, computed analogously to the trastuzumab epitope described above.

### Biological Baselines

We used the Observed Antibody Space (OAS)^7^ and the Structural Antibody Database (SAbDab)^8^ to generate sets of biologically relevant sequences for comparison to our generative models.

For HCDR3: The “Random OAS” baseline was constructed by randomly sampling 50,000 unique HCDR3s from OAS with the only condition being that they had a length of 9-17 amino acid residues (for parity with our model generations). We similarly constructed the “Random OAS-J” baseline by randomly sampling 10,000 unique HCDR3s from OAS from antibodies that have the same J-gene as trastuzumab, while also imposing the same length constraint as the OAS baseline. For the “SAbDab” baseline, we included all 2,395 unique HCDR3s from SAbDab that have lengths of 9-17 residues and were neither trastuzumab nor its variants. For the “Permuted Sequences” baseline, we randomly sampled a subset of 5,000 HCDR3s from the “Random OAS” baseline and randomly shuffled each amino acid sequence via permutation to destroy positional information.

For all three HCDRs (HCDR123): We took an analogous approach to the HCDR3 baselines but restricted it to HCDR1 and HCDR2 sequence lengths of 8 residues (to match the HCDR1 and HCDR2 lengths of trastuzumab). We sampled 50,000 sequences for the “Random OAS” baseline, 10,000 sequences for the “Random OAS-J” baseline, and all 1,572 unique HCDR123s from SAbDab, which fit the length criteria for the “SAbDab” baseline and were neither trastuzumab nor its variants.

For analysis post *in vitro* screening, we removed from consideration any CDRs that contained cysteines to avoid unpaired cysteines and match this sampling constraint in IgMPNN. We present results with and without cysteines (Supplementary Table 4) and find that binding rates are higher among CDRs that do not contain cysteines compared to CDRs that do (*p* < 1e-5 for OAS, *p* = 0.0014 for OAS-J, Fisher’s exact test).

A fraction of each library was also used for ablation studies (results not presented; used for internal benchmarking), so the overall binding rate of the entire libraries (consisting of hundreds of thousands of sequences) may appear lower than the top *k* binding rates reported here.

### Binding Rate of Top **k** Sequences

We defined the binding rate as the fraction of designs in a population of size k which are determined experimentally to bind to the target antigen. To compute the binding rate, we first identified the top k sequences; these are determined via ranking by sequence loss (model log-likelihood) within groups of fixed HCDR3 lengths weighted by the OAS-J length distribution (Supplementary Table 1). Specifically:

- We defined Top_f_(k_f_) binders as the number of binders amongst the top k_f_ sequences (top sequences according to model log-likelihood) with HCDR3 lengths equal to f.
- We let *f_f_* be the frequency of length *f* HCDR3s in OAS-J for *f E {9,10,., 17}*. Then we took *(k_9_, k_l0_,., k_l7_)* such that 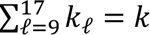 and the quantity 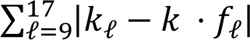 minimized.
- We let Top*(k)* 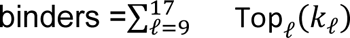 binders.

In some cases, the number of HCDR3s with length 9 or length 10 screened was lower than *k_9_, k_l0_* as computed above. In that case, we considered all such HCDR3s and recomputed *k_ll_, k_l2_,., k_l7_* via the same strategy originally used for *k_9_, k_l0_,., k_l7_*, but with the budget adjusted to the difference between *k* and the number of HCDR3s with length 9 or length 10 and with the OAS-J frequency modified to only consider *f > 10*.

### Comparing Binding Rates

For comparing binding rates between two populations, we ran Fisher’s exact test. Specifically, if population 1 consisted of *b_l_*binders and *n_l_* non-binders and population 2 consisted of *b_2_* binders and *n_2_* non-binders then:

- The binding rates for population 1 and population 2 were given by 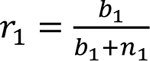, and 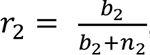 respectively.
- The ratio of population 1’s binding rate to population 2’s binding rate was *<u>^rl^</u>r2*.
- The *p*-value (Fisher’s exact test) corresponding to the binding rates of population 1 and 2 was 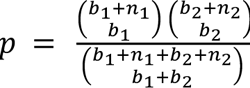

## *In Silico* Structural Modeling

Three-dimensional models of selected *de novo* HCDR3 binders were created in PyMOL^16^ and the Crystallographic Object-Oriented Toolkit (*Coot*)^17^ using the coordinates of the trastuzumab-HER2 complex (PDB:1N8Z). Rosetta’s FastRelax application^18^ was applied using flexible backbone and side-chain degrees of freedom parameters. Prior to the relax procedure, we first idealized all candidate structures using Rosetta’s Idealize protocol to avoid steric clashes and improper geometry. We relaxed the structures using the maximum number of rotamers by passing *-EX1, -EX2, -EX3* and *-EX4* flags at initialization. We also included flags *-packing:repack_only* to disable design, *-no_his_his_pairE*, and *-multi_cool_annealer 10* to set the number of annealing iterations. For ranking of conformations in FastRelax, we used Rosetta’s REF2015 energy function. It is well known that running relax on a structure will often move the backbone a few Angstroms^b^, so we included an additional term containing harmonic distance constraints for all pairs of *C/3* atoms that were either not part of a CDR loop or not within distance 10 Å to any atom in a CDR loop, based on the conformation of the initial structure. These constraints were given weight 1e^-4^. The protocol was run 10 times for each target, and we selected the decoy with the lowest energy in the HCDR3 loop.

### Cloning

Antibody variants were cloned and expressed in Fab format. To produce ACE Assay^TM^ and surface plasmon resonance (SPR) datasets, DNA variants of HCDR3 alone or spanning HCDR1 to HCDR3 were purchased as single-stranded DNA (ssDNA) oligo pools (Twist Bioscience). We spot-checked selected binders by re-purchasing as double stranded DNA eBlocks (Integrated DNA Technologies) or ssDNA oligo pools. Codons were randomly selected from the two most common in *E. coli* B strain^19^ for each residue.

Amplification of the ssDNA oligo pools was carried out by PCR according to Twist Bioscience’s recommendations, except for using Q5 high fidelity DNA polymerase (New England Biolabs) instead of KAPA polymerase. Briefly, 25 µL reactions consisted of 1x Q5 Mastermix, forward and reverse primers, 0.3 µM each, and 10 ng oligo pool. Reactions were initially denatured for 3 min at 95°C, followed by 13 cycles of: 95°C for 20 s; 66°C for 20 s; 72°C for 15 s; and a final extension of 72°C for 1 min. DNA amplification was confirmed by agarose gel electrophoresis, and amplified DNA was subsequently purified (DNA Clean and Concentrate Kit, Zymo Research).

To generate linearized vector, a two-step PCR was carried out to split our plasmid vector carrying the Fab format trastuzumab into two fragments in a manner that provided cloning overlaps of approximately 25 nucleotides (nt) on the 5’ and 3’ ends of the amplified ssDNA oligo pool libraries, or 40 nt on the 5’ and 3’ ends of IDT eBlocks (Integrated DNA Technologies).

Vector linearization reactions were digested with DpnI (New England Biolabs) and purified from a 0.8% agarose gel using the Gel DNA Recovery Kit (Zymo Research) to eliminate parental vector carry through. Cloning reactions consisted of 50 fmol of each purified vector fragment, either 100 fmol PCR-amplified ssDNA oligo pool, or 10 pmol eBlock library inserts and 1x final concentration NEBuilder HiFi DNA Assembly (New England Biolabs). Reactions were incubated at 50°C for 25 min using eBlocks or 2 h using PCR-amplified oligo pools. Assemblies were subsequently purified using the DNA Clean and Concentrate Kit (Zymo Research). DNA concentrations were measured using a NanoDrop OneC (Thermo Scientific).

For high-diversity libraries (HDLs), Transformax EPI300 (Lucigen) *E. coli* was transformed using the MicroPulser Electroporator (BioRad) with the purified assembly reactions and recovered in 1000 µL of SOC medium cultivated at 30°C for 1 h. The cell culture was then grown in 20 mL of Teknova LB Broth with 50 µg/mL Kanamycin at 30°C and 80% humidity with 270 rpm shaking for 18 h. Plasmids were extracted (Plasmid Midi Kit, Zymo Research) and submitted for QC sequencing. Electrocompetent SoluPro™ host strain was transformed with 20 ng of DNA and recovered in 500 µL of SOC medium cultivated at 30° for 1 h.

For low-diversity libraries (LDLs), Absci SoluPro™ host strain was transformed with the purified assembly reactions and grown overnight at 30°C on agar plates containing 50 µg/ml kanamycin and 1% glucose. Colonies were picked for QC analysis prior to cultivation for induction.

### Quality Control Analysis

The quality of high-diversity variant libraries was assessed by deep sequencing. Briefly, library plasmid pools were amplified by PCR across the region of interest and sequenced with 2×150 or 2×300 nt reads using the Illumina MiSeq platform with 20% PhiX. The PCR reaction used 10 nM primer concentration, Q5 2× master mix (New England Biolabs), and 1 ng of input DNA diluted in H2O. Reactions were initially denatured at 98°C for 3 min; followed by 30 cycles of 98°C for 10 s, 59°C for 30 s, 72°C for 15 s; with a final extension of 72°C for 2 min. Sequencing results were analyzed for distribution of mutations, variant representation, library complexity, and recovery of expected sequences. Metrics included coefficient of variation of sequence representation, read share of top 1% most prevalent sequences, and percentage of designed library sequences observed within the library.

The quality of low-diversity variant libraries was assessed by performing rolling circle amplification (Equiphi29, Thermo Fisher Scientific) on 24 colonies and sequencing using the Illumina DNA Prep, Tagmentation Kit (Illumina Inc.). Each colony was analyzed for mutations from reference sequence, presence of multiple variants, misassembly, and matching to a library sequence (Geneious Prime).

### Antibody Expression in SoluProM *E. coli* B Strain

After recovery in SOC medium, high-diversity libraries (HDLs) were grown in 50 mL of Teknova LB Broth with 50 µg/mL Kanamycin at 30°C and 80% humidity with 270 rpm shaking for 24 h. After 24 h, the pre-culture was diluted to OD600 = 1 in 100 mL induction base medium (IBM) (4.5 g/L potassium phosphate monobasic, 13.8 g/L ammonium sulfate, 20.5 g/L yeast extract, 20.5 g/L glycerol, 1.95 g/L citric aAcid) containing inducers and supplements (250 µM arabinose, 50 µg/mL Kanamycin, 8 mM magnesium sulfate, 1 mM propionate, 1X Korz trace metals) and grown for 16 h in a 500 mL baffled flask at 26°C and 80% humidity with 270 rpm shaking. At the end of the 16 h, 250 µL aliquots adjusted to 20% v/v glycerol were stored at −80°C.

After transformation and QC of low-diversity libraries (LDLs), individual colonies were picked into deep-well plates containing 400 µL of Teknova LB Broth 50 µg/mL Kanamycin and incubated at 30°C and 80% humidity with 1000 rpm shaking for 24 h. At the end of the 24 h, 150 µL samples were centrifuged (3300 g, 7 min), supernatant decanted from the pre-culture plate, and cell pellets sent for sequence analysis. Samples of 80 µL from the pre-culture were transferred to 400 µL of IBM containing inducers and supplements as described above. Culture was grown for 16 h at 26°C and 80% humidity with 270 rpm shaking. After 16 h, 150 µL samples were taken and centrifuged (3300 g, 7 min) into pellets with supernatant decanting prior to being stored at −80°C.

### Activity-specific Cell-Enrichment (ACE) Assay**^TM^**

For staining, thawed glycerol stocks from induced cultures were transferred to 0.7 ml matrix tubes (500 µL, OD600 = 2), centrifuged (4000 g, 5 min), and resulting pelleted cells were washed three times with PBS (pH 7.4, 1 mM µL of phosphate buffer by adding prior to fixation 250 µL of 0.6% paraformaldehyde and 0.04% glutaraldehyde in phosphate buffer (32 mM, pH 7.4)). After 40 min incubation on ice, samples were centrifuged (4000 g, 5 min) and pellets were washed three times with PBS (pH 7.4, 1 mM EDTA), resuspended in permeabilization buffer (20 mM Tris, 50 mM glucose, 10 mM EDTA, 5 µg/mL rLysozyme), and incubated for 8 min on ice.

Fixed and permeabilized cells were then centrifuged (4000 g, 5 min) and washed three times with staining buffer (Perkin Elmer AlphaLISA immunoassay buffer, 25 mM HEPES, 0.1% casein, 1 mg/mL dextran-500, 0.5% Triton X-100, 0.05% Kathon).

### Staining

Prior to library staining, the HER2 probe was titrated against the reference strain to determine the 75% effective concentration (EC75). Following cell preparation, the library was resuspended in 500 µL staining buffer containing 100 nM either His/Avi-tagged human HER2 (Acro Biosystems) conjugated to 50 nM streptavidin-AF647 (Invitrogen) or tag-free human HER2 (Acro Biosystems) directly conjugated to AF647 via free amines. Libraries were incubated with the probe overnight (16 h) with end-to-end rotation at 4°C, centrifuged (4000 g, 5 min), and pellets were washed three times with PBS. Pellets were resuspended in 500 µL of staining buffer containing 26.5 nM anti-kappa light chain:BV421 (BioLegend), incubated for 2 h with end-to-end rotation at 4°C prior to centrifugation (4000xg, 5 min), and then washed three times with PBS and resuspended in 200 µL of PBS for sorting.

### Sorting

For the binary ACE Assay^TM^, high-diversity libraries (HDLs) were sorted on FACSymphony S6 (BD Biosciences) instruments. Immediately prior to sorting, 50 µL of stained sample was transferred to a flow tube containing 1 mL PBS + 3 µL propidium iodide. Aggregates, debris, and impermeable cells were removed with singlets, size, and PI+ parent gating, respectively. Cells were then gated to include only those with kappa light chain expression (BV421). A total of three collection gates were set to sample at the high end of the binding range (top 90% of the positive binding signal as determined by a sample matched negative control), the remaining 10% positive binding signal events, and a negative gate containing the events with no binding signal. Libraries were sorted simultaneously on up to four instruments with photomultipliers adjusted to normalize fluorescence intensity, and the collected events were processed independently as technical replicates.

### Gating

Gating was done in the following order (visual presented in Supplementary Fig. 14):

1. Singlets were gated to exclude aggregates from analysis using SP SSC-W vs. SP SSC-A.
2. Permeabilized cells were gated through inclusion of all PI positive events on a PI-A vs. SP SSC-A plot to eliminate non-permeabilized cells and non-cell noise and debris.
3. Cell size was evaluated using SSC-A vs FSC-A to ensure only appropriately sized cells were sorted.
4. Kappa light chain expressing cells were gated using a histogram of BV421 signal and a non-expressing control strain as a negative control.
5. HER2 binding cells were gated using a histogram of AF647 and a non-binding control strain as a negative control. HER2 binding was divided into two separate gates (P1 and P2) for analysis within the ACE Assay^TM^ pipeline.

### Sorted Material Sample Preparation

Cell material from sorted gates was collected in a diluted PBS mixture (VWR) in 1.5 mL tubes (Eppendorf). A sample of the unsorted library material was also processed for QC and ACE Assay^TM^ metric calculations. Post-sort samples were centrifuged (3,800 g) and tube volume was normalized to 20 µL. Amplicons encompassing the HCDR3 or VH region were generated by PCR. The reaction used 10 nM primer concentration, Q5 2× master mix (New England Biolabs) and 20 µL of sorted cell material input suspended in diluted PBS (VWR). Reactions were initially denatured at 98°C for 3 min, followed by 30 cycles of 98°C for 10 s; 59°C for 30 s; 72°C for 15 s; with a final extension of 72°C for 2 min. After amplification, samples were cleaned enzymatically using ExoSAP-IT (Applied Biosystems). Resulting DNA samples were quantified by Qubit fluorometer (Invitrogen), prepped for sequencing with the ThruPLEX DNA-Seq Kit (Takara Bio), normalized and pooled. Pool size was verified via Tapestation 1000 HS and was sequenced on Illumina NextSeq 1000 P2 (2×150 nt or 2×300nt) with 20% PhiX.

### Binary ACE Assay**^TM^** Analysis

Enrichment scores were calculated for individual variants screened by a binary version of the ACE Assay^TM^ using the following procedure:

1. Paired-end reads were merged using Fastp^20^ with quality filtering and base correction in merged regions enabled.
2. Primers were removed from both ends of the merged read using the Cutadapt tool^21^, and reads were discarded where primers were not detected.
3. Unique sequences were tallied to provide raw counts of each variant observed in each sample. Sequences that did not match a designed sequence in the library were discarded.
4. For each sample, proportional abundances were calculated for each variant. Enrichment scores were calculated by dividing the proportional abundance of each variant in a gate by its proportional abundance in the unsorted library sample.

### Surface Plasmon Resonance (SPR)

#### Sample Preparation

Post-induction samples were transferred to 96-well plates (Greiner Bio-One), pelleted and lysed in 50 µL lysis buffer (1X BugBuster protein extraction reagent (Millipore) containing 0.01 KU Benzonase nuclease (Millipore) and 1X Halt Protease inhibitor cocktail (Thermo Scientific)).

Plates were incubated for 15-20 min at 30°C then centrifuged to remove insoluble debris. After lysis, samples were adjusted with 200 µL SPR running buffer (10 mM HEPES, 150 mM NaCl, 3 mM EDTA, 0.01% w/v Tween-20, 0.5 mg/mL BSA) to a final volume of 260 µL and filtered into 96-well plates. Lysed samples were then transferred from 96-well plates to 384-well plates for high-throughput SPR using a Hamilton STAR automated liquid handler. Colonies were prepared in two sets of independent replicates prior to lysis and each replicate was measured in two separate experimental runs. In some instances, single replicates were used, as indicated.

### SPR

High-throughput SPR experiments were conducted on a Carterra LSA SPR instrument using SPR running buffer (10 mM HEPES, 150 mM NaCl, 3 mM EDTA, 0.05% w/v Tween-20, 0.5 mg/mL BSA) and SPR wash buffer (10 mM HEPES, 150 mM NaCl, 3 mM EDTA, 0.05% w/v Tween-20). Carterra LSA SAHC30M chips were functionalized with 20 µg/mL biotinylated antibody capture reagent for 600 s prior to conducting experiments. Lysed samples in 384-well blocks were immobilized onto chip surfaces for 600 s followed by a 60 s washout step for baseline stabilization. Antigen binding was conducted using a 300 s association phase followed by a 900 s dissociation phase. Six leading blanks of SPR running buffer were injected to create a consistent baseline prior to monitoring antigen binding kinetics. After the leading blanks, five concentrations of HER2 extracellular domain (ACRO Biosystems, three-fold serial dilutions from 500 nM) were injected. After antigen injections, the chip was regenerated with two 120 s injections of regeneration buffer (10 mM glycine, pH 2.0). Sample immobilization, blanks, and antigen injections were repeated to produce technical replicate data. In some cases, biological replicates were also run separately for additional replicate data points per clone.

### Sequencing for SPR Libraries (LDLs)

To identify the DNA sequence of individual antibody variants evaluated by SPR, duplicate plates were provided for sequencing. A portion of the pelleted material was transferred into 96 well PCR (Thermo-Fisher) plate via pinner (Fisher Scientific) which contained reagents for performing an initial phase PCR of a two-phase PCR for addition of Illumina adapters and sequencing. The reaction volume was 12.5 µL. During the initial PCR phase, partial Illumina adapters were added to the amplicon via 4 PCR cycles. The second phase PCR added the remaining portion of the Illumina sequencing adapter and the Illumina i5 and i7 sample indices. The initial PCR reaction used 0.45 µM UMI primer concentration, 6.25 µL Q5 2× master mix (New England Biolabs) and PCR-grade H2O. Reactions were initially denatured at 98°C for 3 min, followed by 4 cycles of 98°C for 10 s; 59°C for 30 s; 72°C for 30 s; with a final extension of 72°C for 2 min. Following the initial PCR, 0.5 µM of the secondary sample index primers were added to each reaction tube. Reactions were then denatured at 98°C for 3 min, followed by 29 cycles of 98°C for 10 s; 62°C for 30 s; 72°C for 15 s; with a final extension of 72°C for 2 min.

Reactions were then pooled into a 1.5 mL tube (Eppendorf). Pooled samples were size-selected with a 1x AMPure XP (Beckman Coulter) bead procedure. Resulting DNA samples were quantified by Qubit fluorometer. Pool size was verified via Tapestation 1000 HS and was sequenced on Illumina MiSeq Micro (2×150 nt) for HCDR3 libraries or an Illumina MiSeq Reagent Kit v3 (2×300 nt) for HCDR1-HCDR3 libraries with 20% PhiX.

After sequencing, amplicon reads were merged using Fastp^20^, trimmed by cutadapt^21^ and each unique sequence enumerated. Next, custom R scripts were applied to calculate sequence frequency ratios between the most abundant and second-most abundant sequences in each sample. Levenshtein distance was also calculated between the two sequences. These values were used for downstream filtering to ensure a clonal population was measured by SPR. The most abundant sequence within each sample was compared to the designed sequences and discarded if it did not match any expected sequence. Dominant sequences were then combined with their companion Carterra SPR measurements.

### Binder Identification with the Activity-specific Cell-Enrichment (ACE) Assay^TM^

To determine the success of the ACE Assay^TM^, we included over a thousand controls (SPR-validated binders and non-binders) in the libraries. The binary ACE Assay^TM^ (bACE) produced enrichment scores based on proportional abundances in the specified FACS gates. The *P_l_* and *P_2_* enrichment scores were predictive of binding (Supplementary Fig. 1) based on their separation of the binding and non-binding controls.

To label screened sequences as binders, we set a threshold on the median *P_l_* enrichment score (across three replicates, *R_l_, R_2_, R_3_*) and a separate threshold on the minimum *P_2_* enrichment score (across the same three replicates as *P_l_*). Specifically, given thresholds *t_l_, t_2_*, we labeled a sequence a binder if:

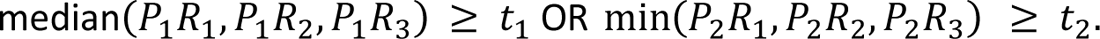

Otherwise, we labeled the sequence a non-binder.

We determined the thresholds t_1_, t_2_ using a grid-search aimed at maximizing F1-score on the controls included in the library. For the HCDR3 library, we found that t_1_ = 3.51, t_2_ = 8.89 achieved the highest F1-score (Supplementary Table 2). For the HCDR123 library, we found that t_1_ = 6.87, t_2_ = 3.05 achieved the highest F1-score (Supplementary Table 3).

### Functional, Developability and Manufacturability Assessment of Final Candidates

#### Cloning expression and purification of high-affinity Fab candidates

High-affinity Fabs were cloned and expressed as described above, and their sequences were verified. Upon seed induction, Fabs were expressed in IBM media in 50 mL shake flask cultures for 16 h (see antibody expression methods above). The cells were pelleted and lysed in lysis buffer (50 mM Tris, 20 mM sodium chloride, 2 mM magnesium chloride, pH 7.5, 2 U/mL Benzonase nuclease (Millipore) and 2400 U/mL rLysozyme Solution (Millipore)). Then, they were incubated at 4°C on a rocking shaker for 1 hr followed by centrifugation at 15,000 RPM. Supernatants were loaded onto CH1-XL affinity resin (Thermo Scientific), washed with PBS and eluted in 10 mM Glycine pH 2.5. Eluted samples were neutralized with 1 M Tris-HCL pH 9. Samples were buffer exchanged into 1X PBS and stored at 4°C.

#### Product Quality of Final Candidates Reformatted as mAbs

All experimental monoclonal antibodies (mAbs) were produced at Wuxi Biologics in a CHO-K1 expression system. Samples were purified via protein A affinity chromatography purification, further purified by size exclusion chromatography (SEC), and kept in a storage buffer.

Concentration of mAbs was determined by A280 using the SoloVPE instrument (Repligen). Suitable protein quality was confirmed by SEC, reduced capillary gel electrophoresis (CGE) or non-reduced CGE (NR-CGE), imaging capillary isoelectric focusing (icIEF), and intact mass spectrometry after deglycosylation. For aggregation determination, SEC was performed using 1X PBS, pH 7.4 as a mobile phase and a 30 cm column TSKgel UP-SW2000 (Tosoh). To evaluate fragmentation, NR-CGE was performed with SDS-MW Analysis Kit (Sciex) together with the SDS sample buffer with iodoacetamide; the separation method was performed using the high-speed setup. When CGE was performed under reducing conditions, 5% (v/v) 2-mercaptoethanol was added to the SDS sample buffer, and the separation method was also performed using high-speed setup. Charge distribution and pI of the candidates were determined by icIEF using the Maurice system (Protein Simple). Samples were prepared by mixing 1:9 (v:v) of 1.0 mg/mL mAb samples and the following master mix: 0.35% (v/v) 1% methylcellulose, 2 M urea, 4% (v/v) Pharmalyte® 3-10 (Cytiva), 0.5% (v/v) pI marker 4.05 (Protein Simple), 0.5% (v/v) pI marker 9.99 (Protein Simple), and 10 mM arginine.

Confirmation of identity and evaluation of highly abundant post-translational modifications were determined by deglycosylated intact mass spectrometry. The samples were deglycosylated using PNGase F (New England Biolabs) overnight and under native conditions as instructed by the manufacturer. Intact mass analysis was performed using a reversed phase HPLC (Agilent) connected to a TripleTOF 6600+ MS System (Sciex). Intact mass data analysis was performed using PMI-Byos software v4.5-53 (Protein Metrics).

One experimental mAb (HCDR3 = ARYRHWYYDYDY) showed < 95% purity by NR-CGE. Data on developabilty, functional assessment, and epitope mapping are released but not presented due to the possibility of manufacturing issues that could affect the side-by-side comparison with the other high-purity candidates.

#### Affinity and Cross-Reactivity SPR

SPR experiments were performed as described above. Human HER2 extracellular domain (Acro Biosystems) was used as an analyte with injections of 3-fold serial dilutions of 500 nM starting concentrations. For affinity determination, sensorgrams were fitted using non-linear regression to a single site binding model. Similar experiments using human HER1 (Acro Biosystems), human HER3 (Acro Biosystems), human HER4 (Acro Biosystems), cynomolgus HER2 (Acro Biosystems), canine HER2 (Sino Biological), and mouse HER2 (Acro Biosystems) were used for binding selectivity assessment. Rovalpitizumab (Twist Biosciences, Genscript) was used as a negative control in these experiments.

#### Cell-Surface Antigen Binding

The cell-surface antigen binding experiment was performed using SK-OV-3 cells (ATCC, HTB-77) in “thaw & use” format. Cells were grown in McCoy’s 5a medium, supplemented with 10% heat-inactivated fetal bovine serum, until 80-90% of confluency was reached. They were detached with EDTA-Trypsin treatment and cryopreserved in a growth medium, supplemented with 5% DMSO at a density of 10^7^cells per mL. On the day of the experiment, cells were thawed by a 2 min incubation in 37°C water bath, centrifuged at 200 g for 5 min at room temperature, and resuspended in McCoy’s 5a medium, supplemented with 10% heat-inactivated fetal bovine serum, at a final cell density of 5×10^5^ cells/mL. A sample of 100 µL of cell suspension was plated per well of a 96-well plate (5×10^4^ cells/well) and incubated at 37°C, 5% CO2 overnight.

Cells were washed with Hank’s balanced salt solution three times, fixed with 3.7% formaldehyde solution, and blocked with 1% BSA. Serially diluted in TBS 1% BSA, antibodies were added to the fixed cell monolayer. After 1 h of incubation, the secondary anti-human Fc antibody-HRP conjugate was added. Binding of the test antibody was assessed by measuring the optical density of the product of TMB oxidation (HRP substrate). To calculate the EC50 values, non-linear regression analysis was performed in GraphPad Prism software (version 10.0.2).

#### Antibody-Dependent Cell Cytotoxicity (ADCC)

Engineered Jurkat cells (Invivogen, jktl-nfat-cd16), stably transformed with a luciferase gene under the control of an NFAT-driven promotor, were used as the effector cells. SK-OV-3 (ATCC, HTB-77) cells were used as the target cells. Target cells were plated into a 96-well plate the night before, as described in the cell-surface antigen binding section. On the day of the experiment, the growth medium was aspirated, and then serially diluted in McCoy’s 5a medium supplemented with 10% fetal bovine serum. The test antibodies were added to the wells, followed by pre-incubation for 2 h. Effector cells were plated in McCoy’s 5a medium supplemented with 10% fetal bovine serum at a final density of 1e^10^ cells/well. The effector to the target cell ratio (E:T) was 2:1. After 18 h of incubation at 37°C, 5% CO2, 20 µL of growth medium was combined with 180 µL of luciferase substrate (Quanti Luc, Invivogen, rep-qlc1) in a solid white 96-well plate. Luminescence intensity was measured in a microplate reader (Envision, Perkin Elmer) immediately. To calculate the EC50 values, non-linear regression analysis was performed in Prism GraphPad software (version 10.0.2).

#### Dynamic Light Scattering and nano Differential Scanning Fluorimetry (DLS/DSF)

Particle size and melting temperature were assessed using a Prometheus Panta (Nanotemper). Antibodies were diluted to 0.5 mg/mL using 1x PBS µLand spun down and loaded into Prometheus High Sensitivity Capillaries (Nanotemper). DLS measurements were taken at 25°C under high sensitivity settings. Thermal denaturation was performed from 25°C to 90°C using a ramp rate of 0.5°C/min. Prometheus Panta software v1.1 (Nanotemper) was used for data analysis.

#### Developability assays

Self-interaction, polyspecificity and hydrophobicity were determined by affinity capture-self interaction nanoparticle spectroscopy (AC-SINS), anti-insulin and anti-DNA ELISA, and hydrophobic interaction chromatography (HIC). AC-SINS was performed as described in Liu et al^22^. ELISAs for polyspecificity anti-DNA and anti-insulin were performed as described in Mouquet et al^23^. HIC was performed as described in Jain et al^24^. Additionally, for assessment of predictive pharmacokinetics of the final candidates, FcRn chromatography was performed as described in Schoch et al^25^. Briakinumab (Leinco Technologies, P/N LT500) and Herceptin (Genentech) were used as positive and negative controls, respectively.

#### Alanine scanning mutagenesis epitope mapping

A wildtype HER2 domain IV fragment that corresponds to residues 510-629 of human HER2 containing a C-terminal His tag was cloned and expressed in SoluProM as described above. Residues within 5 Å of trastuzumab were identified from the crystal structure PDB:1N8Z. Epitope residues were also selected from published data^26^. Alanine variants were cloned and produced in 96-deep well format. Lysis was conducted according to the protocol described above. The reactivity of alanine variants against all experimental mAbs were assessed using high-throughput SPR. The SPR experiments were conducted on a Carterra LSA HT-SPR instrument equipped with a sensor chip (HC30M) at 25 °C. Surface preparation of the HC30M chip was performed using standard amine-coupling reagents. The chip was activated for 600 s with a freshly prepared solution of 0.1 M 1-ethyl-3-(3-dimethylaminopropyl)carbodiimide (EDC) (Thermo Scientific), 0.1 M N-hydroxysulfosuccinimide (Sulfo-NHS) (Thermo Scientific), and 0.1 M MES buffer, pH 5.5 (Carterra). To capture the His-tagged alanine variants, an anti-His monoclonal antibody (Sino Biological) was coupled on the activated chip surface for 900 s at 200 µg/mL diluted into 10 mM sodium acetate, pH 4.5 (Carterra). Unreactive chip surface areas were blocked with 1 M Ethanolamine, pH 8.5 (Carterra) for 600 s.

Twelve replicates of each alanine variant were immobilized onto the pre-functionalized chip surface for 1200 s followed by a 60 s washout step for baseline stabilization. The experimental mAb assessment was conducted using a kinetics method with a 180 s association phase followed by a 180 s dissociation phase. For each experimental antibody injection, three leading blanks were introduced to create a consistent baseline prior to assessing the reactivity of alanine variants against each antibody. After the leading blanks, a single concentration of each antibody at 200 nM was injected into the instrument. After each antibody injection cycle, the chip was regenerated with two 120 s injections of regeneration buffer (10 mM glycine, pH 2.0).

## Data Availability

Open-source data on the anti-HER2 binders and non-binders and the measured binding affinities for all binders are made available at^c^. The data on functionality, developability, and manufacturability for the high-affinity binders that were further characterized are also made available. Data from the ACE Assay^TM^ are withheld until formal publication.

## Code Availability

Code is withheld until formal publication.

## Statistics

Throughout the main text and methods section, we provide details of the statistics utilized. Included in this section is a summary of such methods:

- Two-sided Fisher’s exact tests are used to compare binding rates in Table 1 and Supplementary Table 1. Details on computation are described in the “Comparing Binding Rates” section above. We consider a significance level a = 0.05 and apply Bonferroni corrections with 12 hypothesis tests in Table 1 and 30 hypothesis tests in Supplementary Table 1. Upper bounds on *p* values or precise values are presented in the table captions. All precise values can be computed using the data within the tables and using the data in Supplementary Table 4 for biological baseline populations.
- One-sided Mann-Whitney U tests are used to compare enrichments between binders and non-binders in Supplementary Fig. 1C, 1E. Scipy^d^ (v1.7.3) was used for computation.
- Pearson R and Spearman p are computed along with their corresponding *p* values using Scipy (v.1.7.3). These values are presented in Supplementary Tables 7 and 9, with n = 9 and n = 12, respectively.
- Error bars representing SD are included in Supplementary Fig. 9B, 12B and 12C.
- Confusion matrices comparing binder/non-binder labeling by the ACE Assay^TM^ and surface plasmon resonance (SPR) are presented in Supplementary Tables 2-3. Accuracy, precision, recall, and F1-score are presented as well.
- Discrete histograms are presented in Fig. 2D and 2E and Supplementary Fig. 3A, 3B, 3C, and 3D. The captions include summary statistics, namely the minimum, maximum, median, mean, and SD of the distributions.
- Box plots are presented in Supplementary Fig. 1C, 1E to compare enrichments between binders and non-binders. For these box plots, the center line represents the median, the box limits represent the upper and lower quartiles, the whiskers represent 1.5x the interquartile range, and the points represent the outliers.

## Supplementary Information

### 1. Supplementary Figures

**Supplementary Fig. 1.**
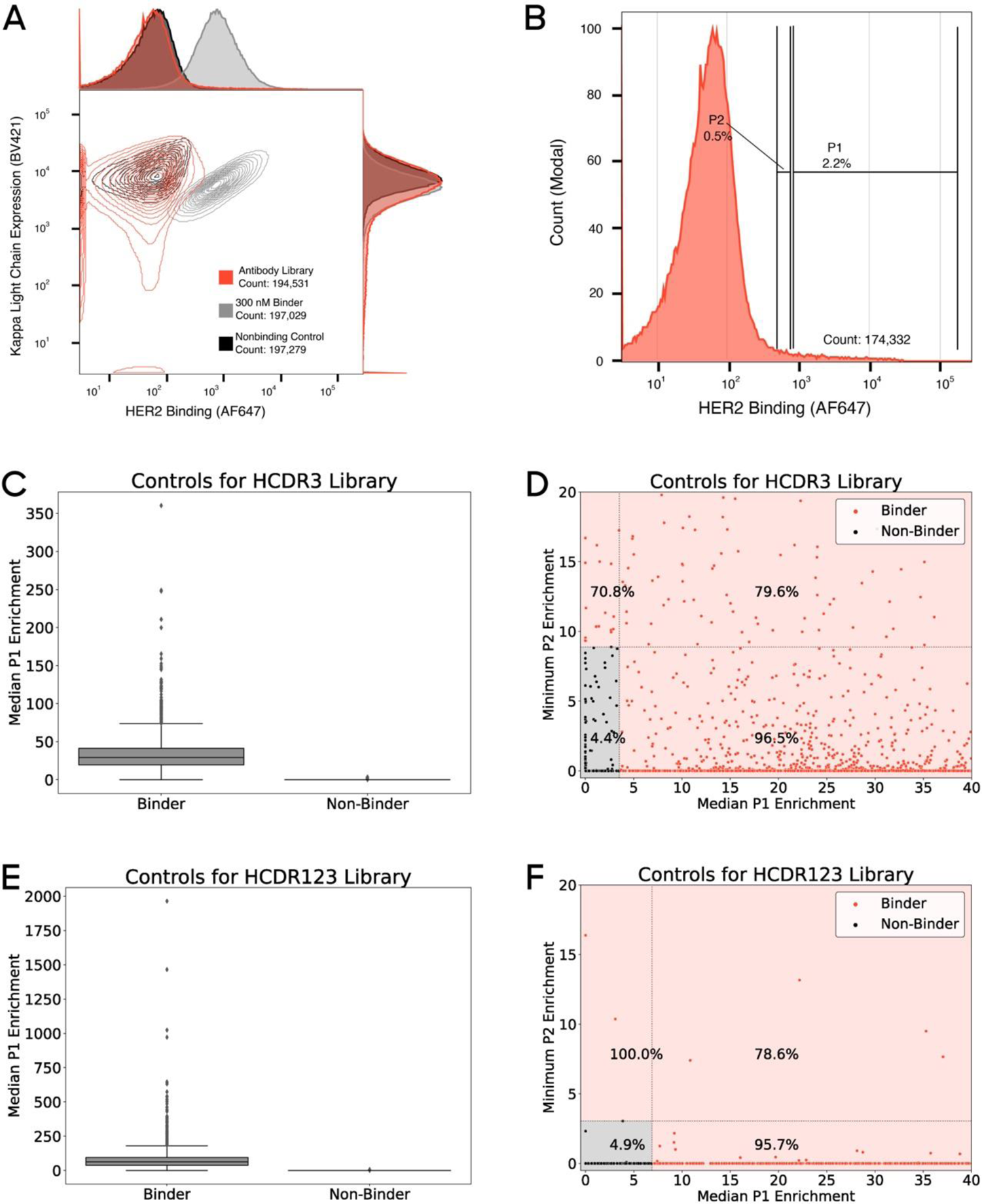
Example of ACE assay and identification of binders from enrichment scores. **(A)** Representative overlay of dot plots showing antigen binding (x-axis, AF647) and antibody expression (y-axis, BV421) for the antibody library (red), low affinity binders (grey), and non-binding controls (black). **(B)** Representative FACS gating strategy used to isolate likely binders from antibody libraries. Gates were set based on non-binding controls. Binders to HER2 are classified in the ACE assay based on median P1 enrichment and minimum P2 enrichment across three replicates (Methods). **(C)** Box plots showing the distribution of median P1 enrichment scores for HCDR3 controls separated by binders and non-binders (center line, median; box limits, upper and lower quartiles; whiskers, 1.5x interquartile range; points, outliers). Binders (n=2203) have statistically significantly higher average median P1 enrichment than non-binders (n=2592) [one-sided Mann-Whitney U test, U = 5509390.5, *p* < 1 x 10^-10^]. **(D)** Plot of HCDR3 controls showing median P1 enrichment scores and minimum P2 enrichment scores. Sequences in the bottom left quadrant (shaded black) are labeled as non-binders, whereas sequences in the other three quadrants (shaded orange) are labeled as binders. The percentage of sequences in each quadrant that are true binders, according to SPR, is shown. Axes truncated to enable better visualization. **(E)** Box plots showing the distribution of median P1 enrichment scores for HCDR123 controls separated by binders and non-binders (center line, median; box limits, upper and lower quartiles; whiskers, 1.5x interquartile range; points, outliers). Binders (n=2270) have statistically significantly higher average median P1 enrichment than non-binders (n=3198) [one-sided Mann-Whitney U test, U = 7003789.5, *p* < 1 x 10^-10^]. **(F)** Plot of HCDR123 controls showing median P1 enrichment scores and minimum P2 enrichment scores. Sequences in the bottom left quadrant (shaded black) are labeled as non-binders, whereas sequences in the other three quadrants (shaded orange) are labeled as binders. The percentage of sequences in each quadrant that are true binders, according to SPR, is shown. Axes truncated to enable better visualization.

**Supplementary Fig. 2.**
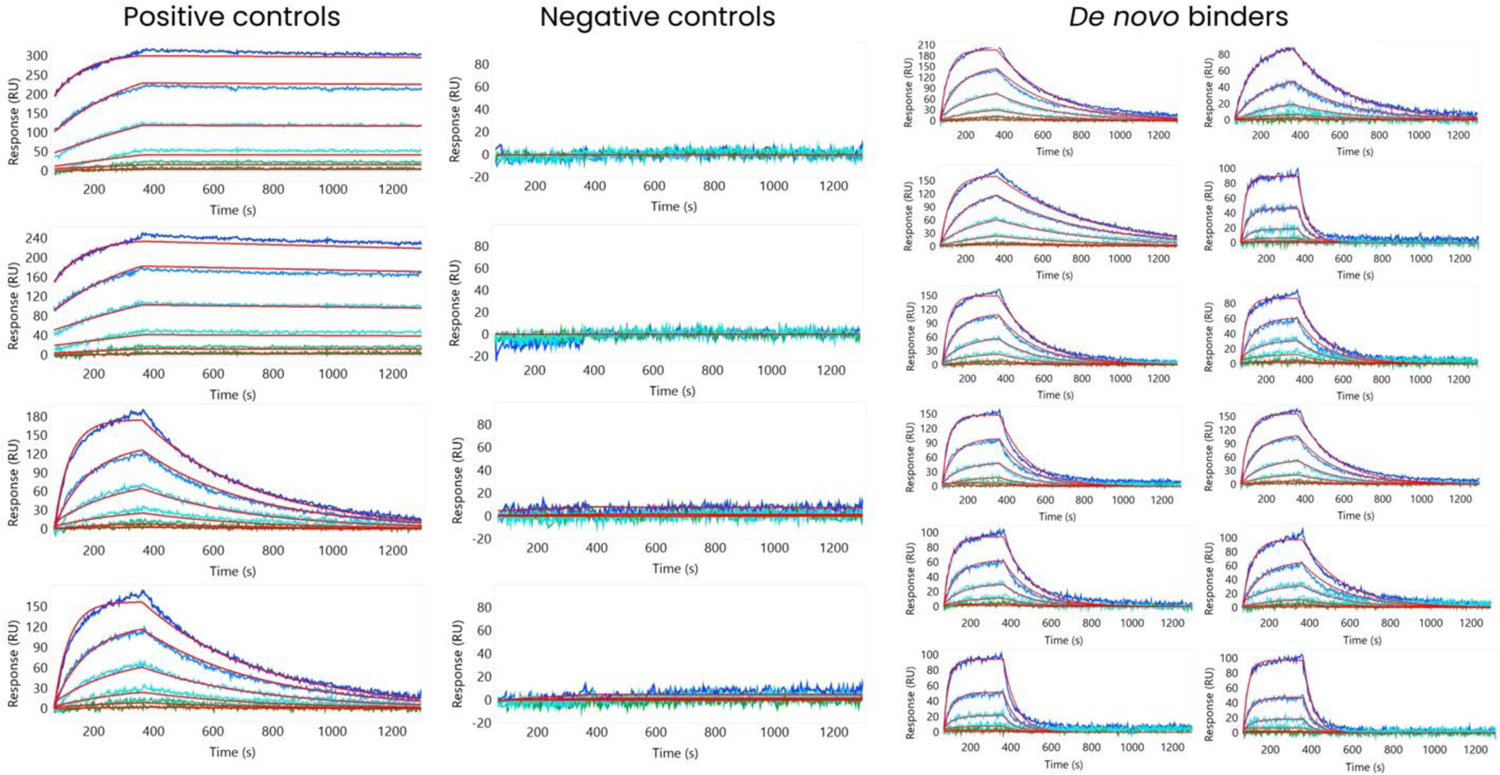
Sensorgram examples of high-throughput SPR workflow for identifying *de novo* binders. (Left) Two positive controls (binders) are shown, each with two replicates. The top-most two sensorgrams show a high-affinity binder; the bottom-most two show a low-affinity binder). **(Center)** Two negative controls (non-binders) are shown, each with two replicates. **(Right)** Two replicates are shown for each of six *de novo* binders. Each row represents one binder.

**Supplementary Fig. 3.**
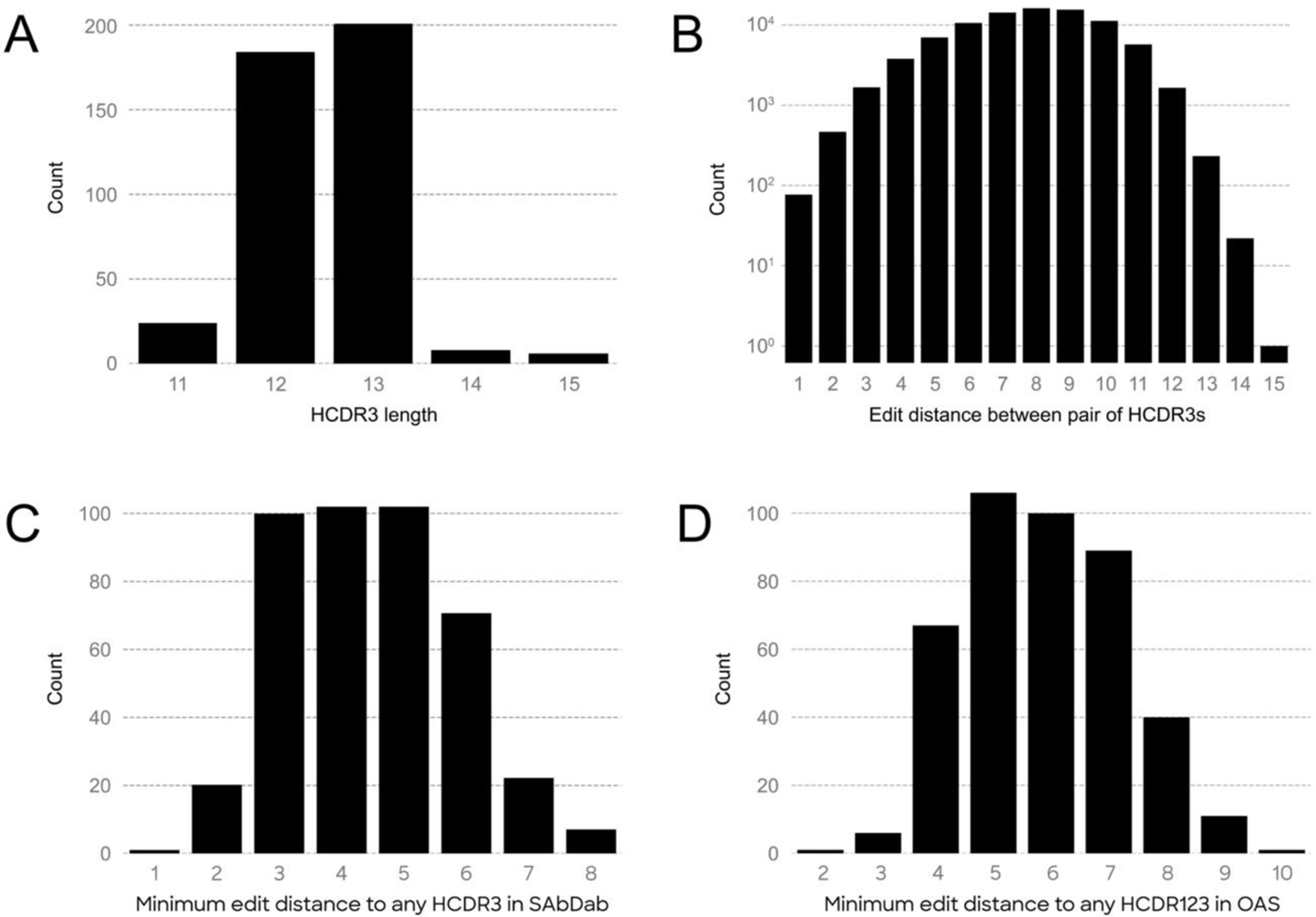
Length and diversity characteristics of *de novo* designed HCDR3 binders. **(A)** Distribution of HCDR3 lengths for zero-shot-designed binders to HER2 (minimum of 11, maximum of 15, median of 13, mean of 12.5 ± 0.69 SD). **(B)** Distribution, on log scale, of pairwise edit distances of zero-shot-designed anti-HER2 HCDR3 binders (minimum of 1, maximum of 15, median of 8, mean of 7.7 ± 2.1 SD). (**C**) Distribution of minimum edit distance to HCDR3s in SAbDab for zero-shot-designed anti-HER2 HCDR3 binders (minimum of 1, maximum of 8, median of 4, mean of 4.46 ± 1.37 SD) (**D**) Distribution of minimum edit distance to HCDR123s in OAS for zero-shot-designed anti-HER2 HCDR3 binders (minimum of 2, maximum of 10, median of 6, mean of 5.87 ± 1.38 SD).

**Supplementary Fig. 4.**
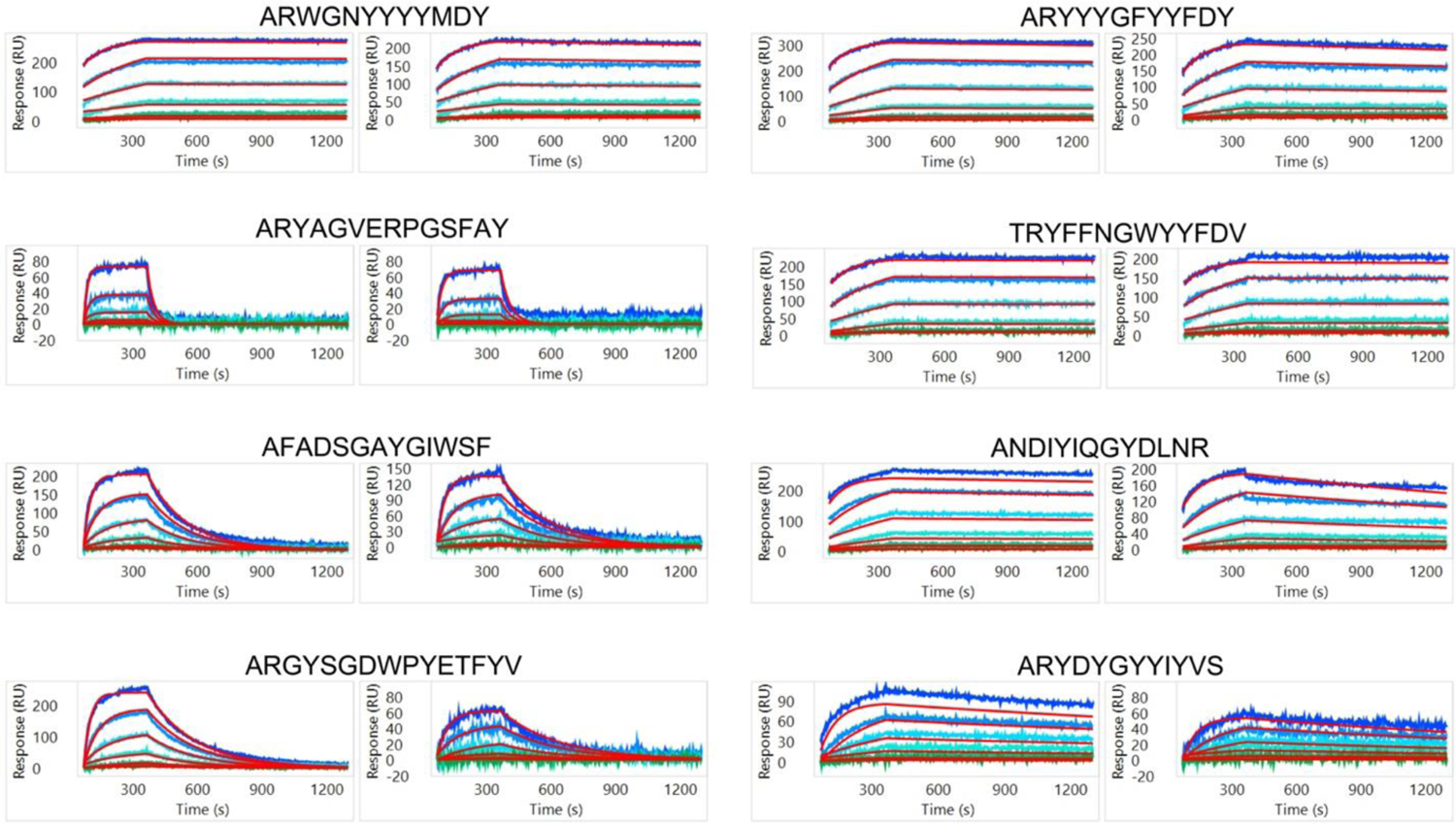
Sensorgrams of eight selected *de novo* HER2 binders for structural modeling. Each set of SPR sensorgrams represents two replicates of each HCDR3 design (sequences on top of each set of sensorgrams).

**Supplementary Fig. 5.**
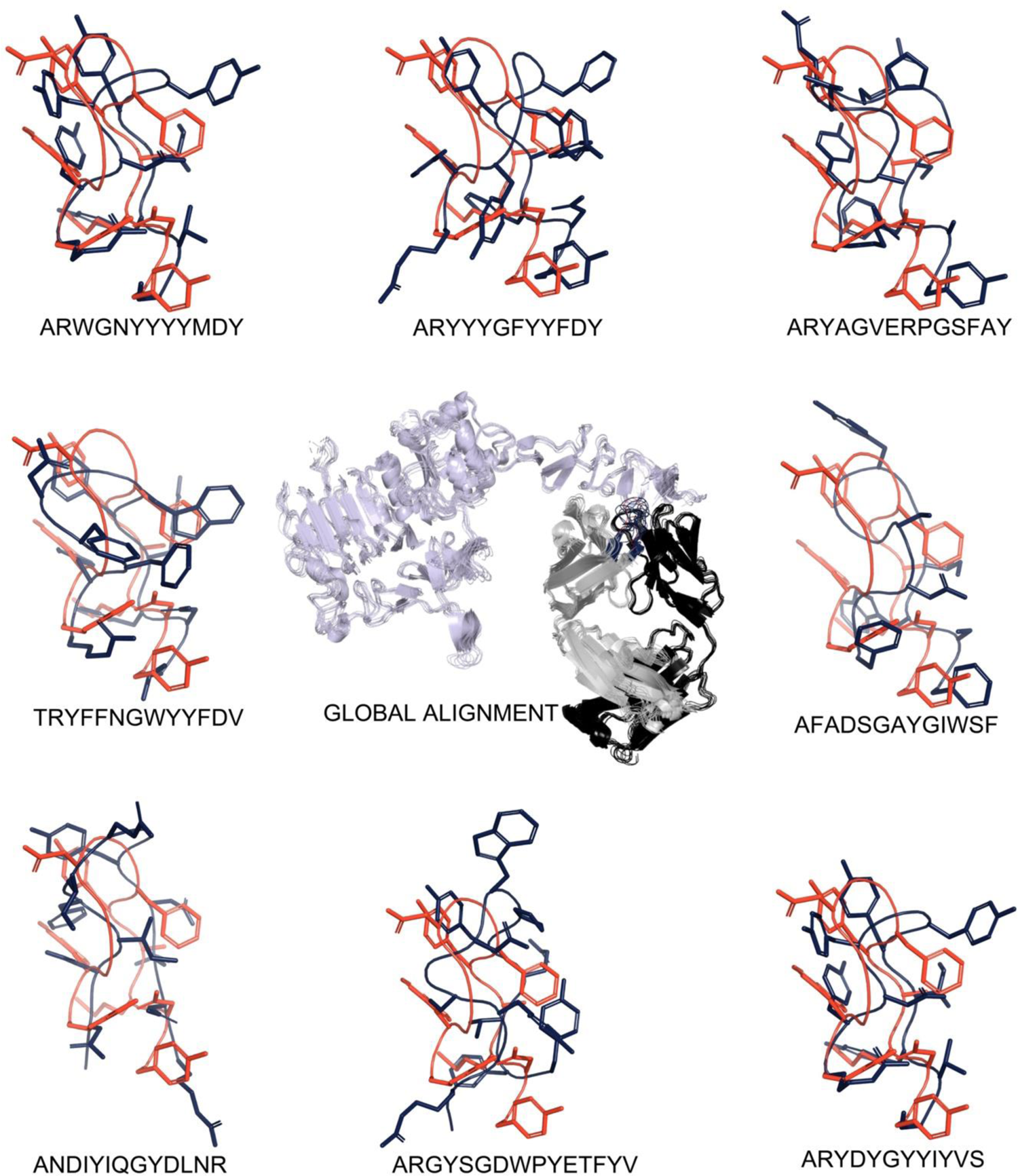
Conformational flexibility of *de novo* designed HCDR3s. Alignment of eight selected *de novo* HER2 binders with trastuzumab-HER2 complex (PDB:1N8Z^1^) shows small overall differences in the antigen (lavender), the heavy chain (gray) and the light chain (dark gray) structures in the overall global alignments (RMSD of 1.1-6.8 Å). Large conformational differences are observed in the *de novo* HCDR3 regions when compared to the conformation of trastuzumab’s HCDR3. The trastuzumab HCDR3 loop is colored red and the *de novo* HCDR3 loops are colored blue. The side chains of the HCDR3 are shown as sticks and the main chain as loops.

**Supplementary Fig. 6.**
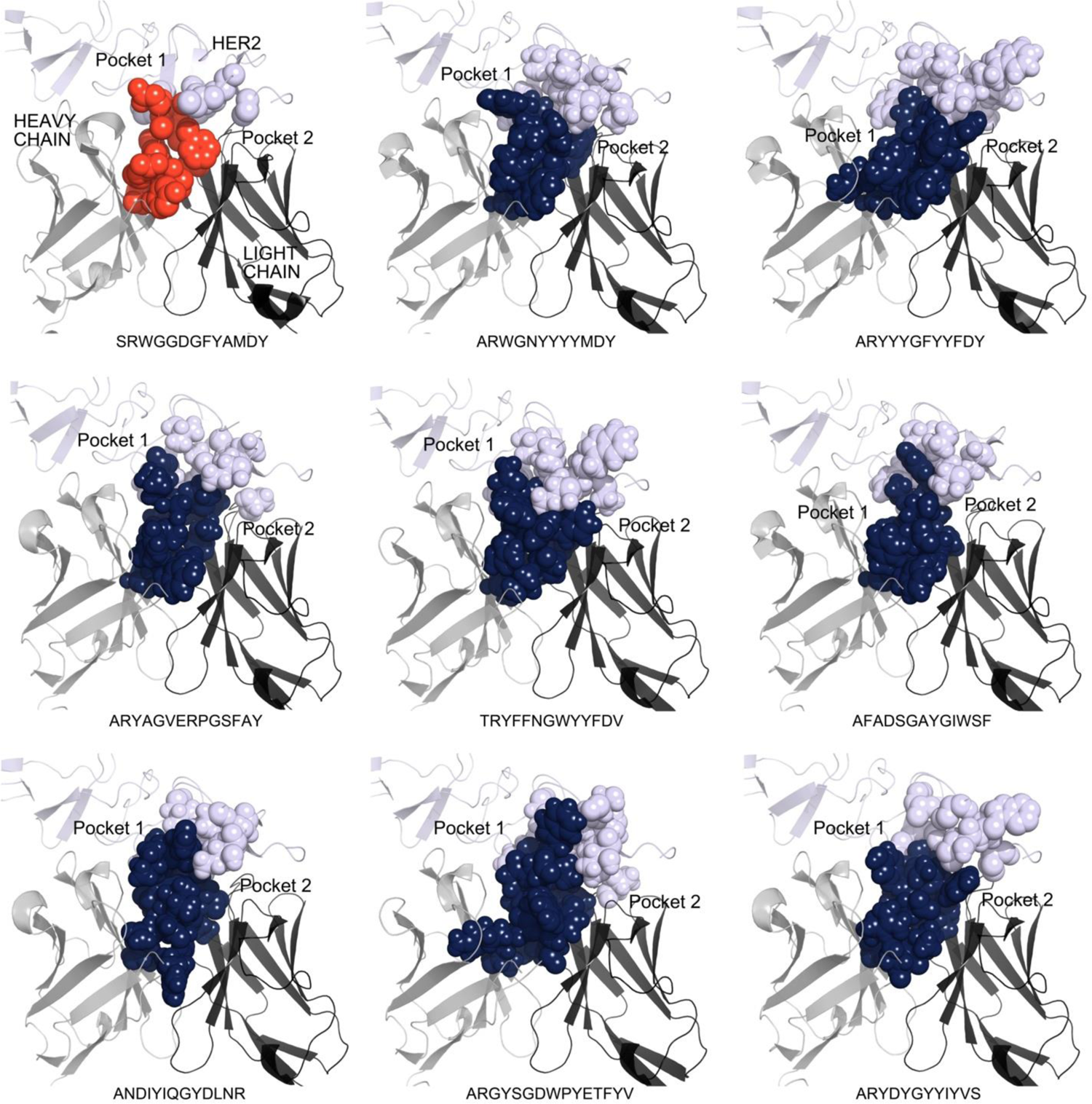
Space-filling representation of HCDR3 loops interacting with epitope residues. Epitope residues were selected using a 5 Å cutoff between the HCDR3 loop and the HER2 domain IV epitope (computed across all atoms). The trastuzumab HCDR3 loop is colored red (top left) and the *de novo* HCDR3 loops are colored blue. Two distinct epitope pockets (Pocket 1 and Pocket 2) that differentially interact with the residues of each HCDR3 can be defined. The interacting surfaces between the HCDR3 and the epitope vary based on HCDR3 sequence and conformation.

**Supplementary Fig. 7.**
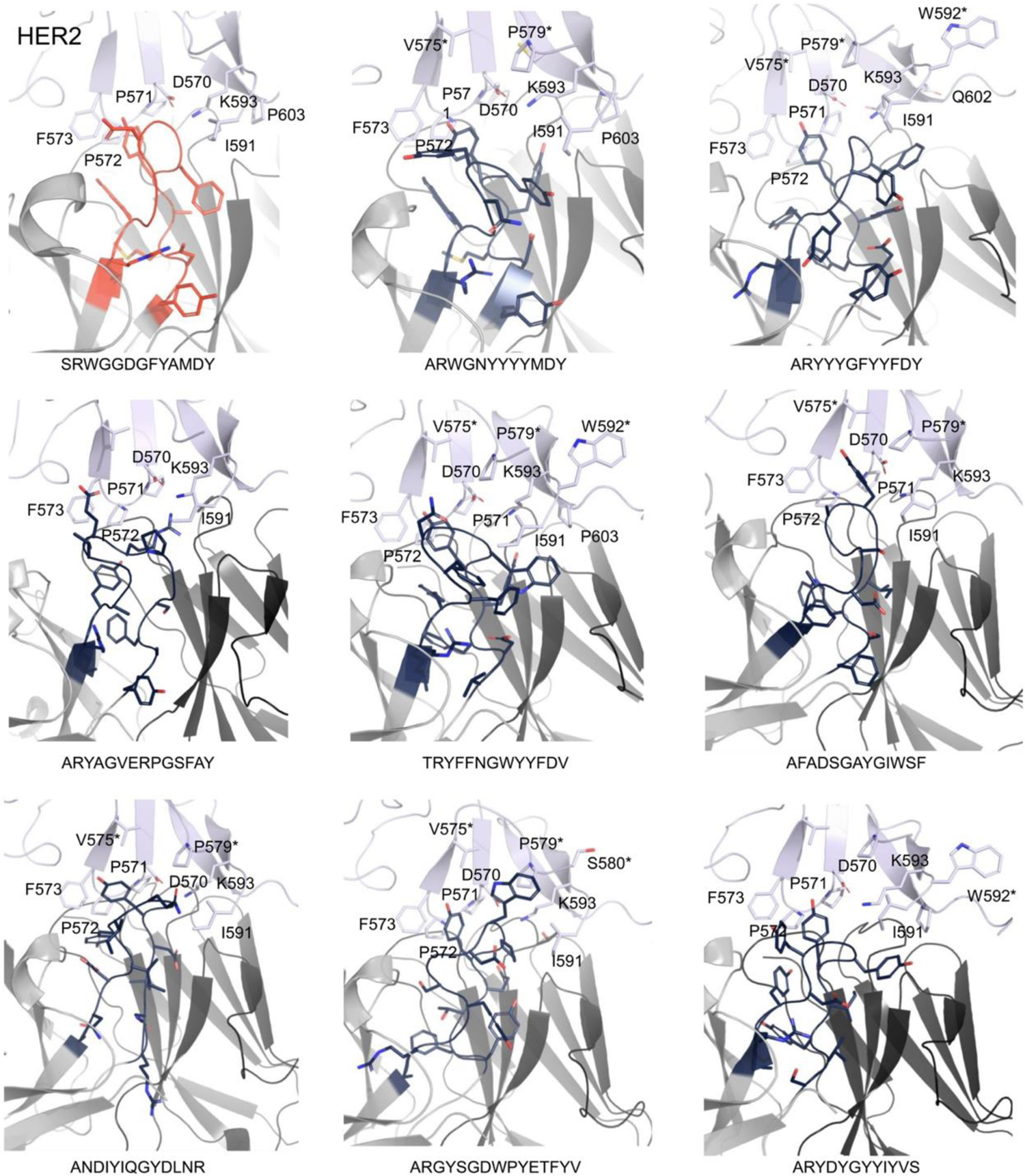
Stick representations of HCDR3-epitope interfaces. Epitope residues were selected using a 5 Å cutoff between HCDR3 and epitope residues (computed across all atoms). The trastuzumab HCDR3 loop is colored red (top left) and the *de novo* HCDR3 loops are colored blue. Epitope residues are labeled according to crystal structure PDB:1N8Z. An *\** denotes novel epitope residues that interact with the *de novo* HCDR3 complexes which are not observed in the trastuzumab-HER2 complex.

**Supplementary Fig. 8.**
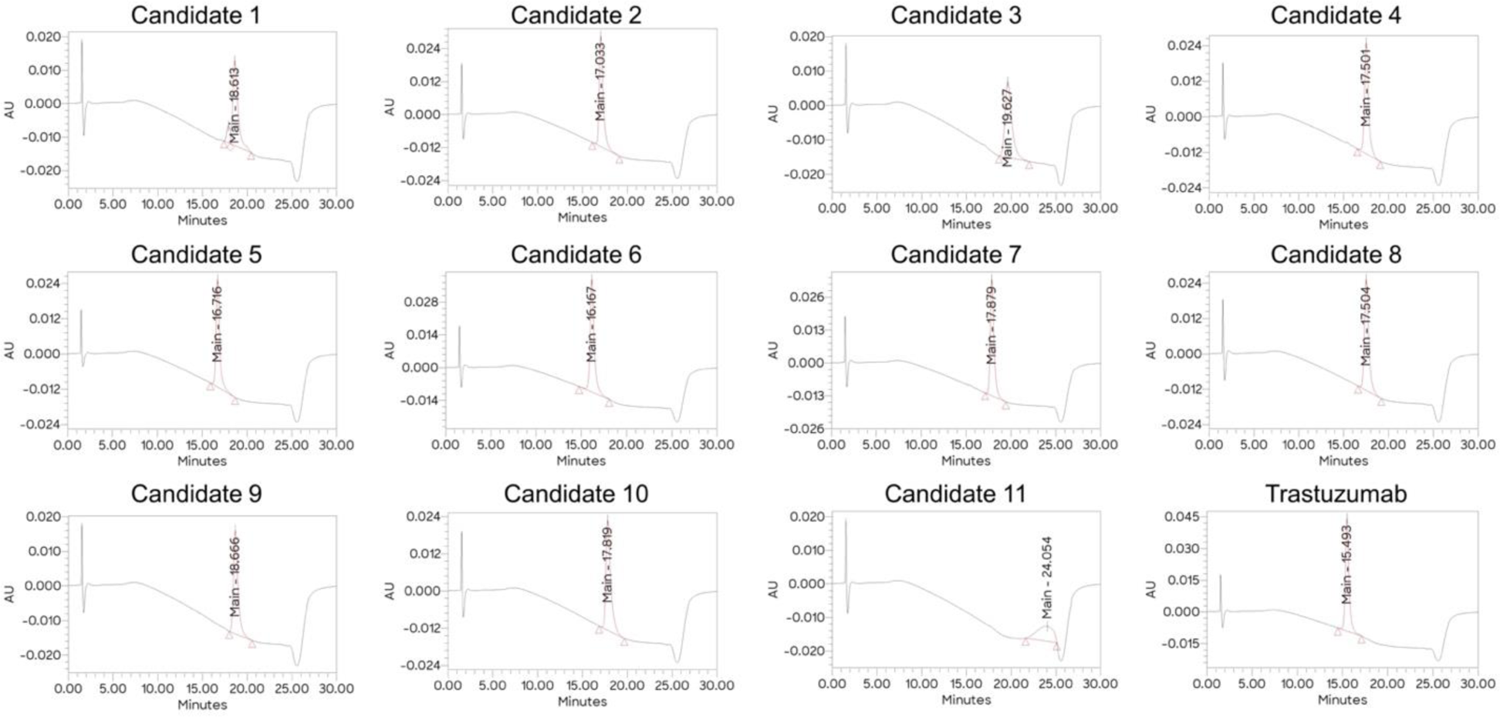
Developability of 11 *de novo* designed antibodies by hydrophobic interaction chromatography (HIC) for protein surface hydrophobicity assessment. Trastuzumab material was analyzed as a sample, whereas Herceptin was used as standard for the relative retention time determination of all candidates.

**Supplementary Fig. 9.**
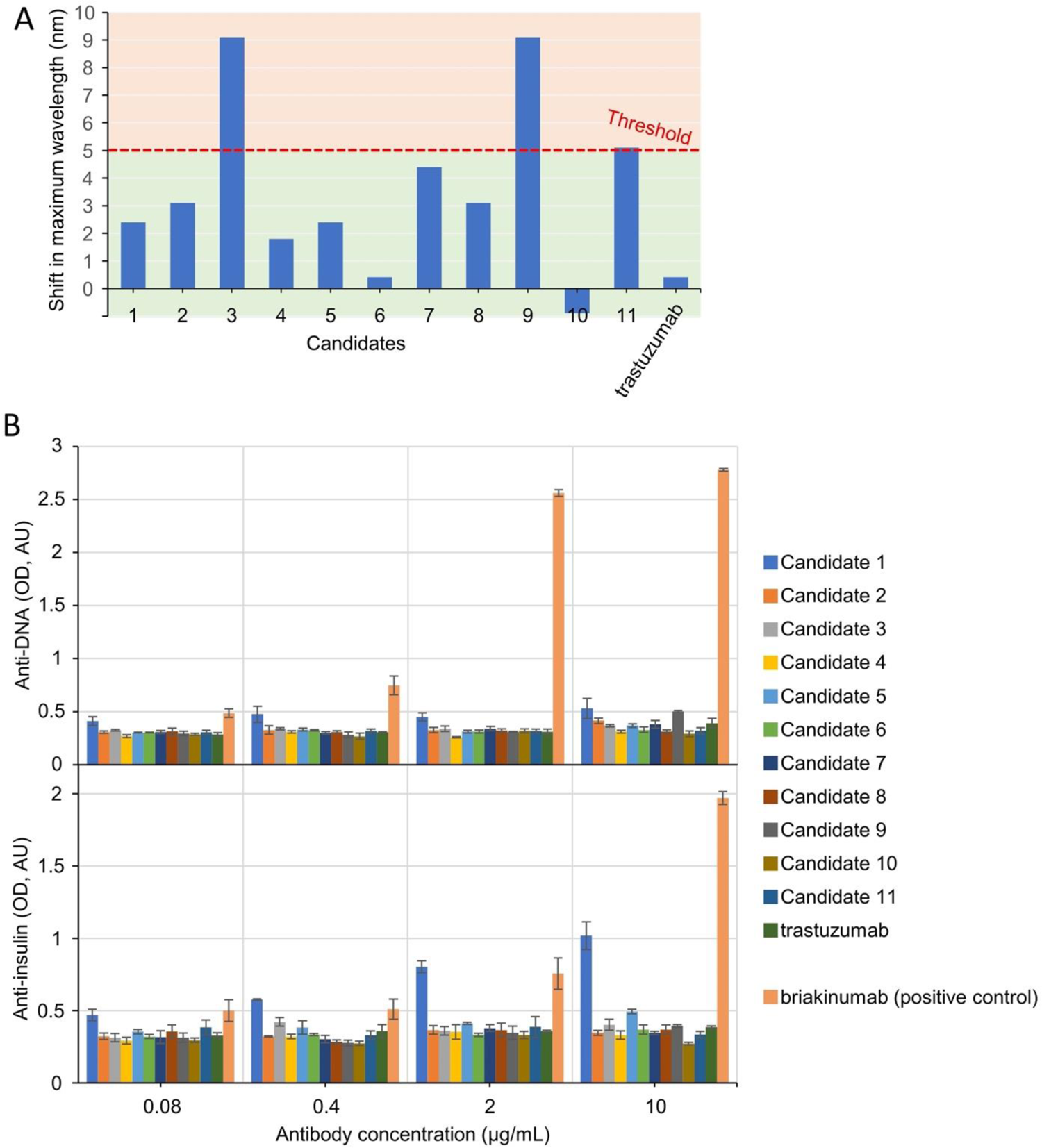
Developability of 11 *de novo* designed antibodies for self-interaction and polyspecificity assessments. Candidates were evaluated by **(A)** AC-SINS for determination of self-interaction and **(B)** anti-DNA (top) or anti-insulin (bottom) ELISA for determination of polyspecificity. The threshold for acceptable AC-SINS is: 5 5nm maximum wavelength shift. Error bars represent SD.

**Supplementary Fig. 10.**
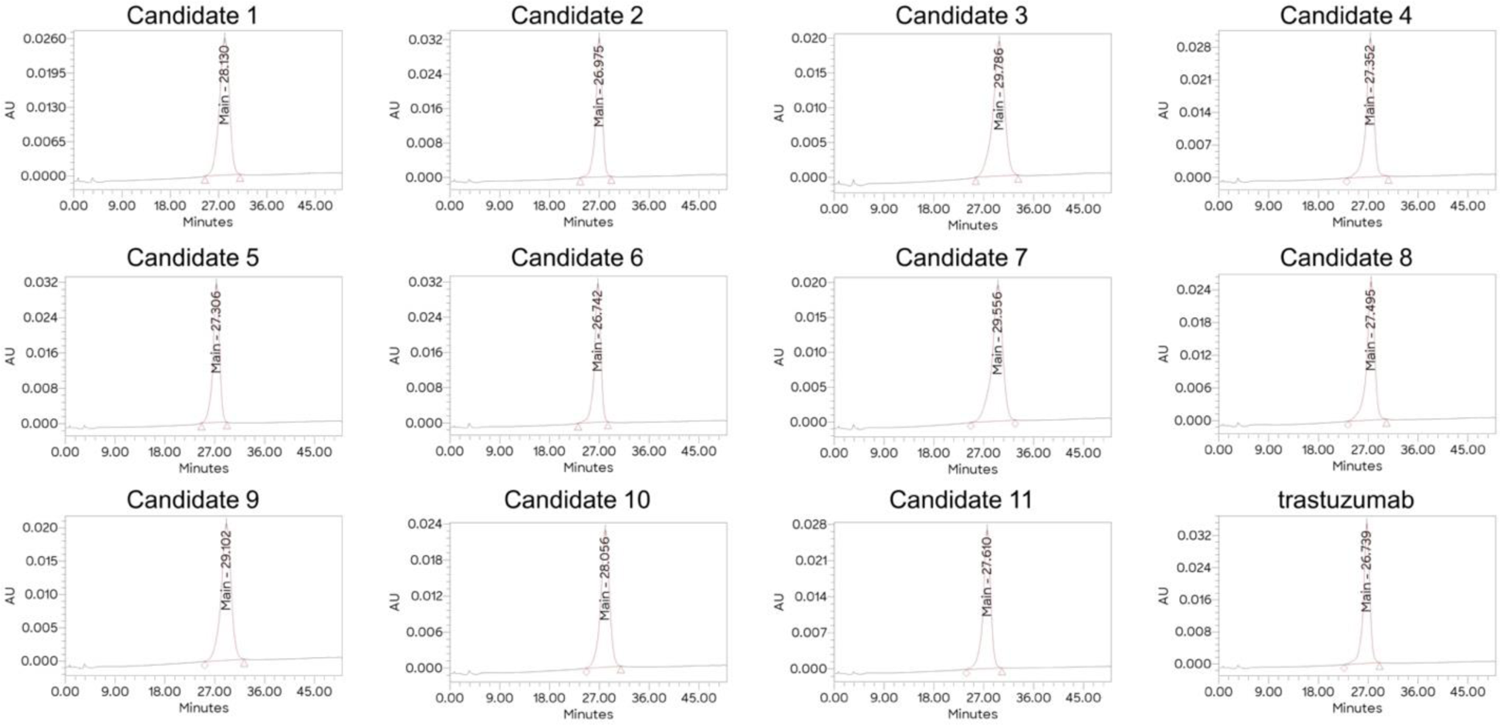
The effects of de novo designed CDRs on FcRn binding for 11 candidate antibodies determined by FcRn chromatography. Batch control trastuzumab material was analyzed as a sample, whereas Herceptin was used as standard for the relative retention time determination of samples.

**Supplementary Fig. 11.**
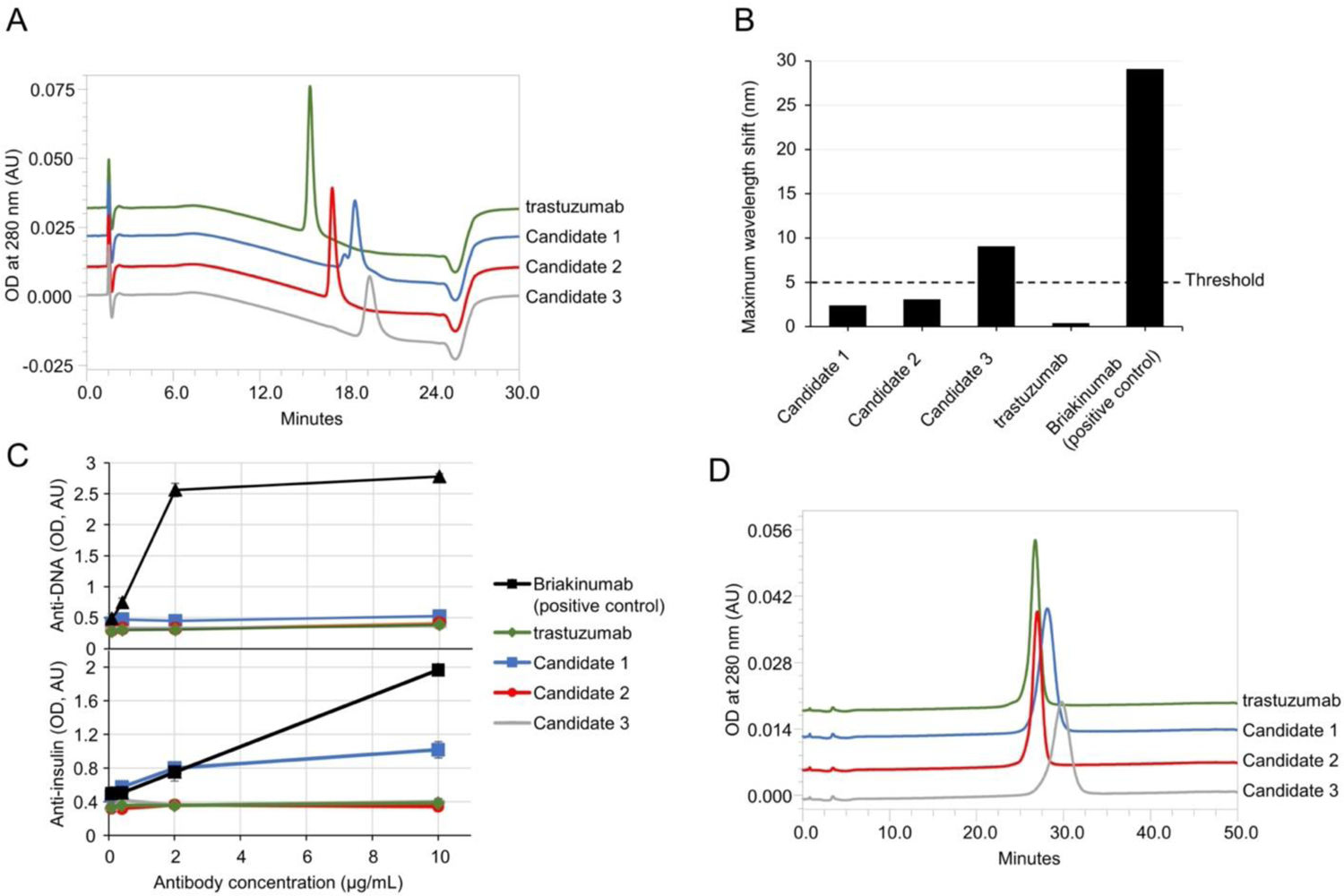
Hydrophobicity, self-interaction, polyspecificity, and FcRn chromatography of *de novo* designed candidates 1-3. Results for **(A)** hydrophobic interaction chromatography (HIC), **(B)** AC-SINS, **(C)** anti-DNA (top) or anti-insulin (bottom) ELISAs, and **(D)** FcRn chromatography are presented side-by-side for Candidates 1-3. Raw data obtained for the assay control briakinumab are shown in selected assays for comparison.

**Supplementary Fig. 12.**
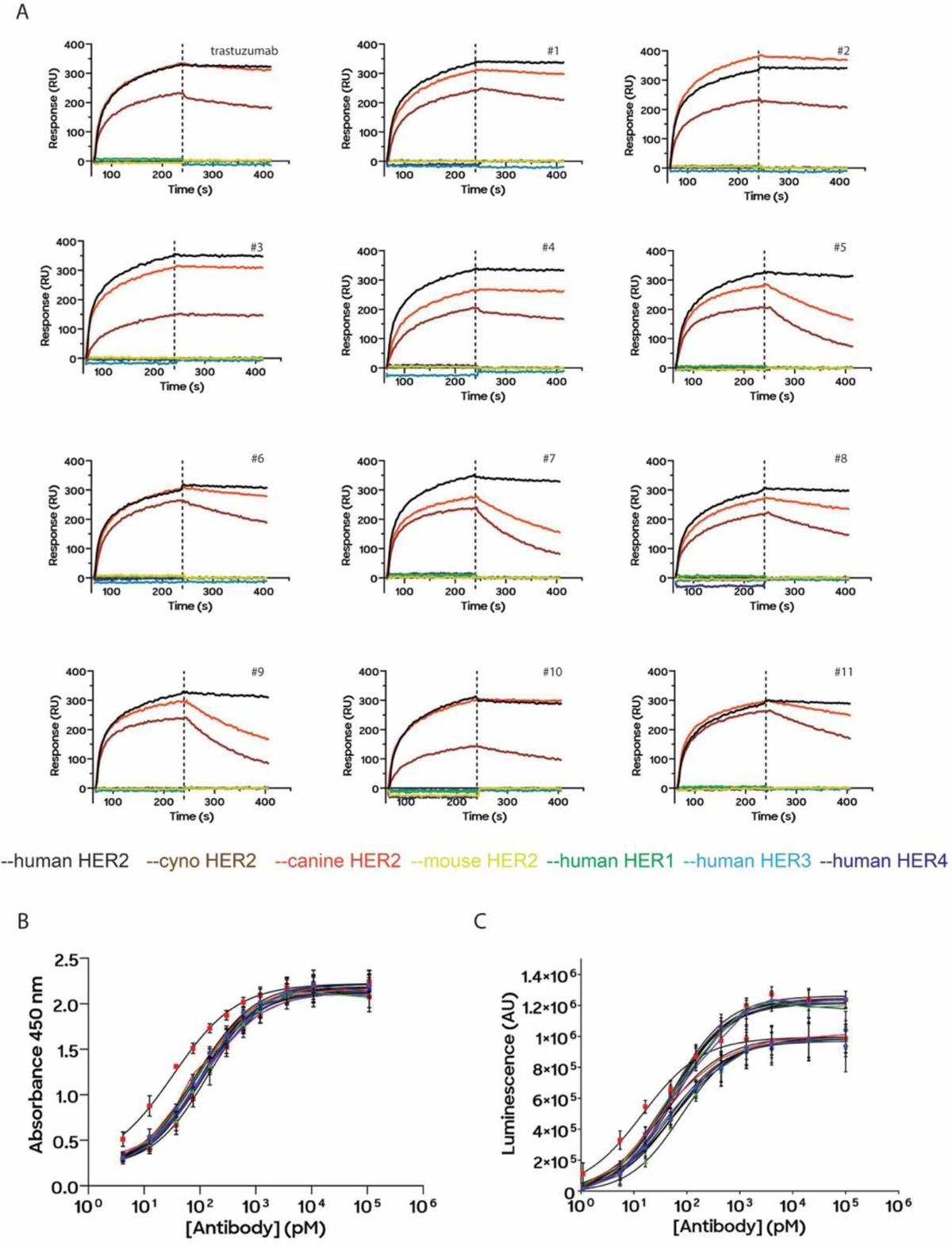
Functional assessment of 11 *de novo* designed antibodies as monoclonal antibodies. **(A)** Cross-reactivity profiles of mAbs. SPR sensorgrams for mAbs binding to a panel of HER2 homologs. Each sensorgram is colored according to figure legend in the trastuzumab plot. All samples were collected in at least 6 single point replicates with 500 nM of each HER2 homolog as analyte. **(B)** Cell-surface binding assay for measuring mAbs binding to HER2^+^ cell line. HRP absorbances as a function of antibody amount was measured over 5 orders of magnitude. **(C)** Antibody-dependent cellular cytotoxicity (ADCC) of Jurkat-Lucia-CD16-NFAT-Luc reporter cell line against HER2^+^ cell line. The activity of secretable luciferase was measured in supernatant. The luciferase activity is proportional to the degree of CD16a engagement. Each value represents the average of triplicate experiments. All titrations are fitted to the hill equation. Error bars represent SD.

**Supplementary Fig. 13.**
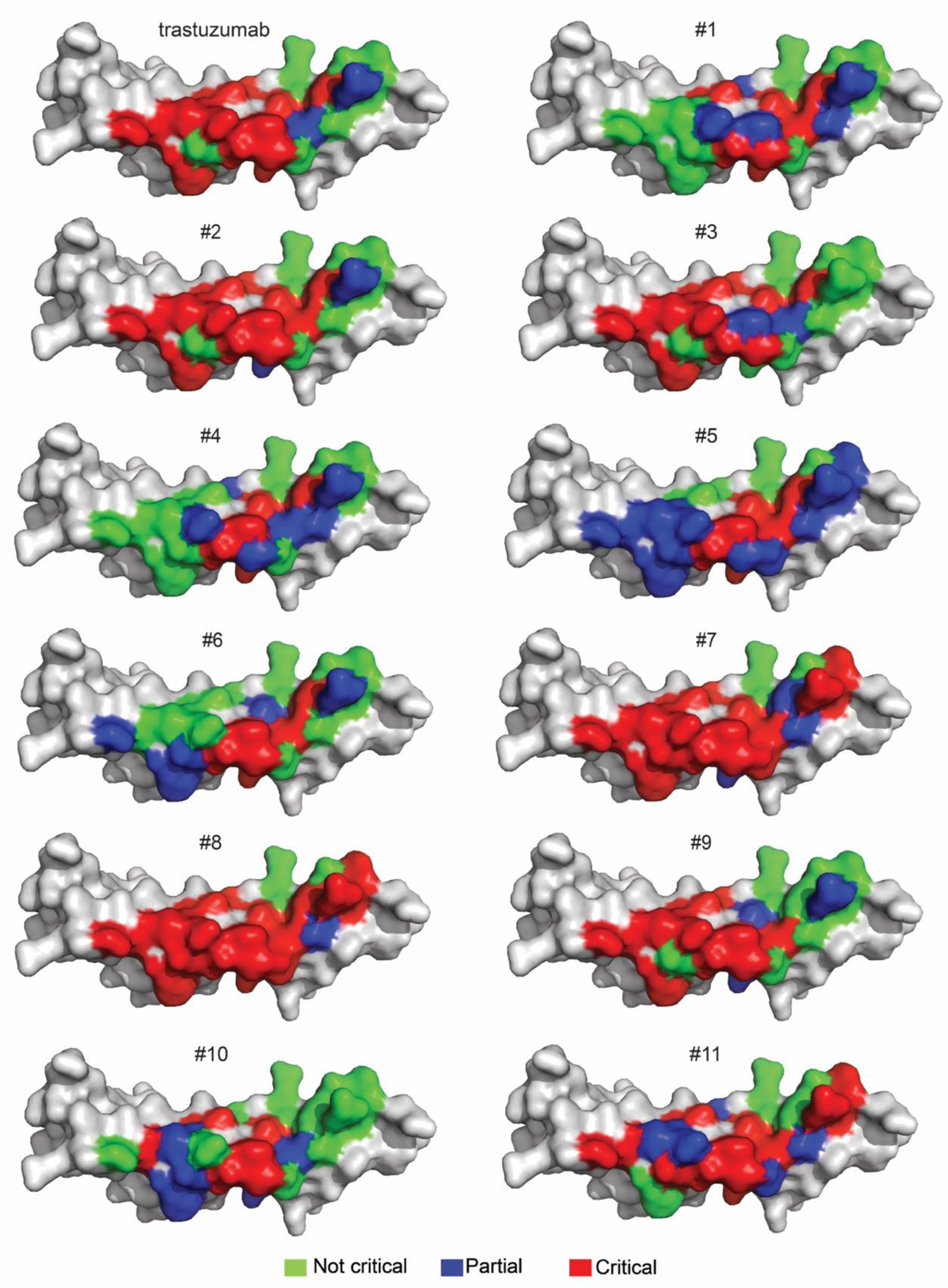
Surface representation of HER2 domain IV alanine scan epitope maps. Residues are colored according to their positions in the trastuzumab-HER2 structure (PDB:1N8Z). Residues that are denoted as not critical have no effect on binding when mutated to alanine. Partially critical residues preserve binding but show greater than 10-fold reduction in binding when mutated to alanine. Critical residues cause a complete loss of binding by SPR when mutated to alanine.

**Supplementary Fig. 14.**
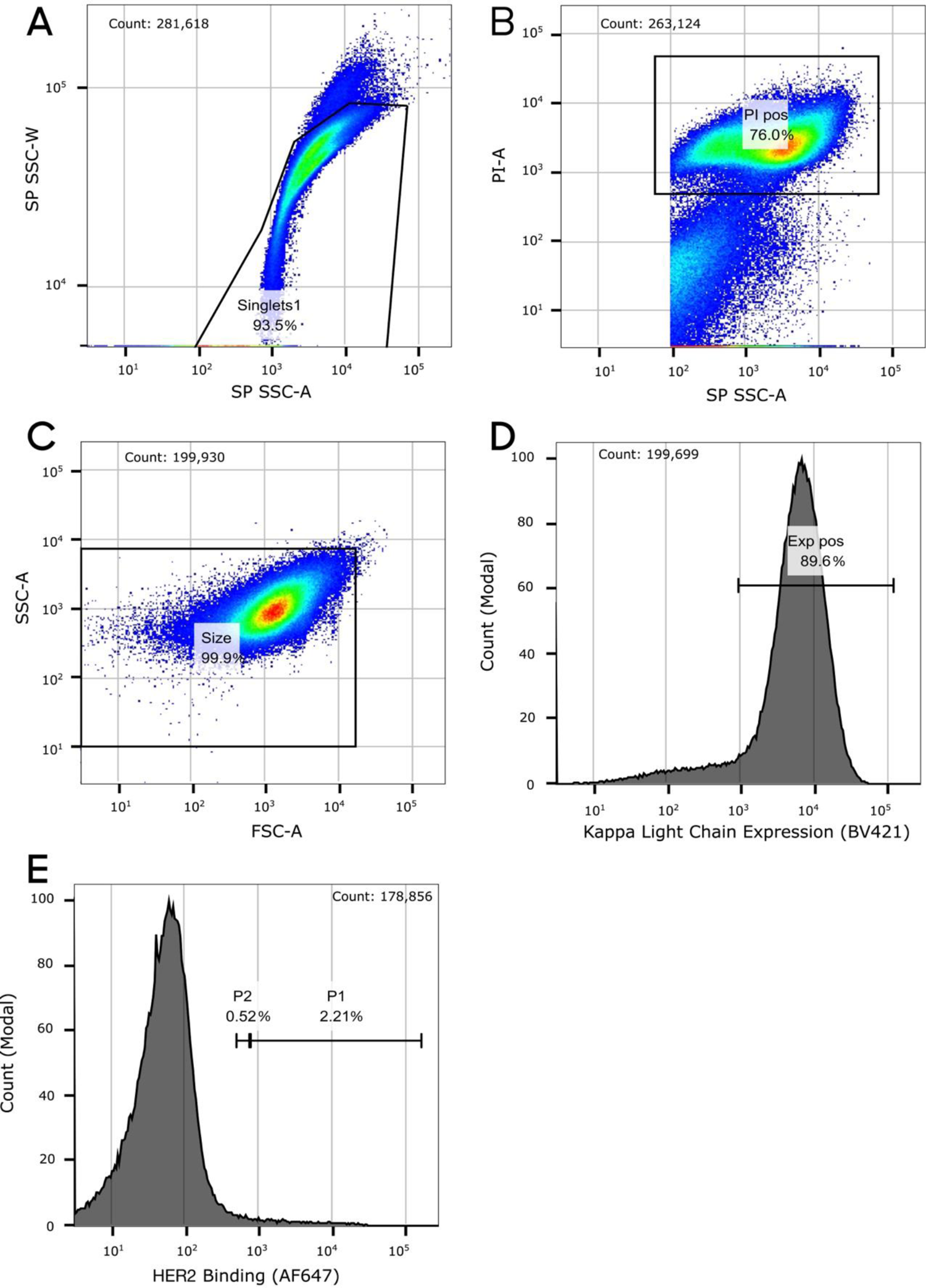
Gating strategy for FACS. Cells were gated by **A)** singlets, **B)** PI staining, **C)** size, **D)** kappa light chain expression, and **E)** HER2 binding. Gates on all fluorescent parameters were set based on negative controls.

### 2. Supplementary Tables

**Supplementary Table 1.**
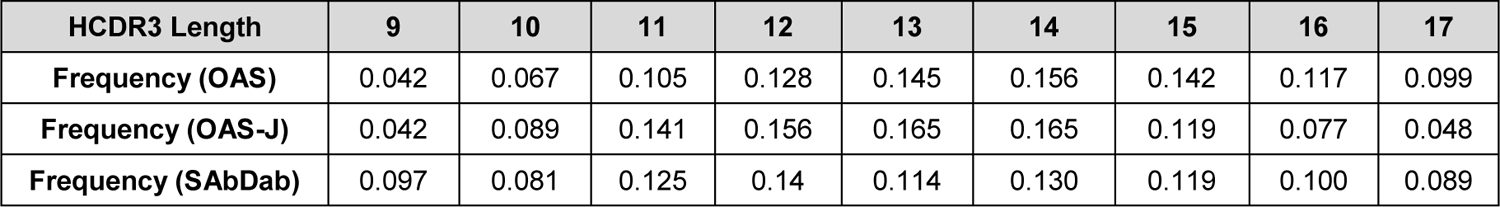
OAS, OAS-J, and SAbDab HCDR3 length distributions. Frequency of HCDR3s appearing in OAS with lengths of 9-17 amino acid residues are shown. OAS-J frequency adds the condition that HCDR3s belong to an antibody with the J gene of trastuzumab. Frequency of unique HCDR3s in SAbDab with lengths of 9-17 residues.

| HCDR3 Length | 9 | 10 | 11 | 12 | 13 | 14 | 15 | 16 | 17 |
| --- | --- | --- | --- | --- | --- | --- | --- | --- | --- |
| Frequency (OAS) | 0.042 | 0.067 | 0.105 | 0.128 | 0.145 | 0.156 | 0.142 | 0.117 | 0.099 |
| Frequency (OAS-J) | 0.042 | 0.089 | 0.141 | 0.156 | 0.165 | 0.165 | 0.119 | 0.077 | 0.048 |
| Frequency (SAbDab) | 0.097 | 0.081 | 0.125 | 0.14 | 0.114 | 0.130 | 0.119 | 0.100 | 0.089 |

**Supplementary Table 2.** ACE Performance on HCDR3 Controls. Confusion matrix for HCDR3 controls indicating binding as measured by SPR and binding as measured by ACE (top). Accuracy, precision, recall, and F1-score are shown (bottom).

|  |  | ACE |  |  |
| --- | --- | --- | --- | --- |
|  |  | Binder | Non-Binder | Total |
| SPR | Binder | 2089 | 114 | 2203 |
|  | Non-Binder | 101 | 2491 | 2592 |
|  | Total | 2190 | 2605 | 4795 |

| Accuracy | Precision | Recall | F1 |
| --- | --- | --- | --- |
| 95.52% | 95.39% | 94.83% | 0.9511 |

**Supplementary Table 3.** ACE Performance on HCDR123 Controls. Confusion matrix for HCDR123 controls indicating binding as measured by SPR and binding as measured by ACE (top). Accuracy, precision, recall, and F1-score are shown (bottom).

|  |  | ACE |  |  |
| --- | --- | --- | --- | --- |
|  |  | Binder | Non-Binder | Total |
| SPR | Binder | 2109 | 161 | 2270 |
|  | Non-Binder | 96 | 3102 | 3198 |
|  | Total | 2205 | 3263 | 5468 |

| Accuracy | Precision | Recall | F1 |
| --- | --- | --- | --- |
| 95.3% | 95.65% | 92.91% | 0.9426 |

**Supplementary Table 4.** Breakdown of sequences experimentally tested for biological baseline populations. Number of binders, total number of sequences, and binding rates for HCDR3 and HCDR123 across different biological baselines. Permuted sequences were not assessed for HCDR123, which is indicated with N/A values. The table on top shows baseline populations after filtering out CDRs that contained cysteines and the table on bottom shows entire baseline populations.

| Population (Cysteines filtered) | # HCDR3 Binders | # HCDR3s Total | HCDR3 Binding Rate (%) | # HCDR123 Binders | # HCDR123s Total | HCDR123 Binding Rate (%) |
| --- | --- | --- | --- | --- | --- | --- |
| OAS | 1,278 | 46,212 | 2.77 | 77 | 44,782 | 0.17 |
| OAS-J | 508 | 9,385 | 5.41 | 30 | 9,253 | 0.32 |
| SAbDab | 74 | 2,271 | 3.16 | 1 | 1,477 | 0.07 |
| Permuted Sequences | 13 | 4,495 | 0.29 | N/A | N/A | N/A |

| Population (Not filtered) | # HCDR3 Binders | # HCDR3s Total | HCDR3 Binding Rate (%) | # HCDR123 Binders | # HCDR123s Total | HCDR123 Binding Rate (%) |
| --- | --- | --- | --- | --- | --- | --- |
| OAS | 1,334 | 49,766 | 2.68 | 79 | 49,346 | 0.16 |
| OAS-J | 521 | 9,921 | 5.25 | 32 | 9,862 | 0.32 |
| SAbDab | 75 | 2,374 | 3.16 | 1 | 1,553 | 0.06 |
| Permuted Sequences | 16 | 4,868 | 0.33 | N/A | N/A | N/A |

**Supplementary Table 5.** *De novo* designs achieve high, calibrated binding rates that outperform biological baselines. The top 100, 1,000, and 10,000 sequences by model likelihood are sampled and tested experimentally. As the number of selected sequences increases, the binding rate decreases. This indicates that the model’s likelihood is calibrated with binding. N/A indicates that fewer than 10,000 sequences were sampled for experimental testing. * Indicates 2-sided Fisher’s exact *p* < 0.001 against each of the biological baselines (OAS, OAS-J, and SAbDab), expect for HCDR3 (k=100) vs. OAS-J with *p* = 0.0021. ° Indicates 2-sided Fisher’s exact *p* < 2 x 10^-4^ against each of the “Incorrect Antigen” populations (rat HER2, HER3, VEGF) except for HCDR3 (k=100) human HER2 vs. rat HER2 and HER3 at *p* = 0.0068 and *p* = 0.0025, respectively, and HCDR123 (k=1,000) human HER2 vs. rat HER2 at *p* = 0.0078.

| Method | HCDR3 Binding Rate (%) |  |  | HCDR123 Binding Rate (%) |  |  |
| --- | --- | --- | --- | --- | --- | --- |
| | $k = 100$ | $k = 1,000$ | $k = 10,000$ | $k = 100$ | $k = 1,000$ | $k = 10,000$ |
| Human HER2 | 13*° | 10.6*° | 6.99*° | 3* | 1.8*° | 1.33* |
| Rat HER2 | 3 | 3.3 | N/A | 0 | 0.6 | N/A |
| Human HER3 | 3 | 2.4 | N/A | 0 | 0.2 | N/A |
| Human VEGF | 2 | 2.5 | N/A | 0 | 0 | N/A |

**Supplementary Table 6.** Binding rates of AI-designed *de novo* designs by HCDR3 length. Top *k* binding rates (i.e., the percentage of the top *k* sequences that were experimentally identified as binders) categorized by HCDR3 length for HCDR3 designs (*k* = 10, 50, 100 for each length) and HCDR123 designs (*k* = 500, 1000 for each length). The baseline binding rates are also presented among the entire population, categorized by HCDR3 length for OAS, OAS-J, and SAbDab.

| Population |  | HCDR3 Length (Binding Rate (%)) |  |  |  |  |  |  |  |  |
| --- | --- | --- | --- | --- | --- | --- | --- | --- | --- | --- |
|  |  | 9 | 10 | 11 | 12 | 13 | 14 | 15 | 16 | 17 |
| HCDR3 | k = 10 | **90** | 20 | 0 | 0 | 10 | 0 | **50** | 0 | 0 |
|  | k = 50 | 54 | **22** | 0 | 6 | 14 | 0 | 24 | 0 | 0 |
|  | k = 100 | 53 | 18 | 1 | 8 | **20** | 1 | 17 | 0 | **2** |
|  | OAS | 14.39 | 5.77 | 3.46 | 4.64 | 2.60 | 1.38 | 0.59 | 0.45 | 0.50 |
|  | OAS-J | 15.67 | 4.97 | **5.77** | **10.80** | 5.94 | **3.05** | 1.94 | **1.17** | 1.54 |
|  | SAbDab | 9.65 | 3.68 | 3.42 | 5.01 | 3.88 | 1.68 | 0.74 | 0.5 | 0 |
| HCDR123 | k = 500 | 1.4 | **10.4** | **1.2** | 0.6 | **0.6** | **0.4** | **0.6** | **0.2** | 0.2 |
|  | k = 1000 | **2.4** | 9.2 | 0.9 | .4 | 0.4 | **0.4** | 0.4 | **0.2** | 0.1 |

**Supplementary Table 7.** Properties of diverse HCDR3 candidates selected for 3D structural modeling. We selected HCDR3 candidates based on affinity, length, and edit distances (ED) to trastuzumab (top). We computed RMSD values over all main chain and side chain atoms from the alignment of HCDR3 residues, excluding all other atoms from calculations. We calculated grand average of hydropathy values (GRAVY) for HCDR3 residues by averaging the hydropathy values of each residue and dividing by sequence length^2^. We computed the Pearson and Spearman correlations between affinity and all other properties (bottom). No significant correlations are seen between affinity (*-log_lO_(K_D_) (M)*) and any of interface area, RMSD, or hydropathy.

| HCDR3 (Length) [ED] | $-\log_{10}(K_D)$ ($M$) | Interface ($\text{\AA}^2$) | RMSD ($\text{\AA}$) | Hydropathy |
| --- | --- | --- | --- | --- |
| SRWGGDGFYAMDY (13) [0] | 8.71 | 771 | 0.000 | -0.81 |
| ARWGNYYYYMDY (12) [6] | 8.77 | 739 | 2.435 | -1.30 |
| ARYYYGFYYFDY (12) [7] | 8.92 | 819 | 2.832 | -0.73 |
| ARYAGVERPGSFAY (14) [11] | 6.24 | 764 | 1.107 | -0.42 |
| TRYFFNGWYYFDV (13) [9] | 9.03 | 843 | 1.974 | -0.37 |
| AFADSGAYGIWSF (13) [12] | 7.00 | 824 | 5.738 | 0.57 |
| ANDIYIQGYDLNR (13) [12] | 8.40 | 833 | 5.506 | -0.80 |
| ARGYSGDWPYETFYV (15) [10] | 7.01 | 863 | 6.767 | -0.76 |
| ARYDYGYYIYVS (12) [10] | 8.02 | 718 | 3.032 | -0.43 |

| Metric | Pearson vs. Affinity |  | Spearman vs. Affinity |  |
| --- | --- | --- | --- | --- |
| | R | p | $\rho$ | p |
| Length | -0.67 | 0.048 | -0.55 | 0.126 |
| Edit distance | -0.54 | 0.132 | -0.67 | 0.048 |
| Interface ($\text{\AA}^2$) | -0.04 | 0.915 | 0.1 | 0.797 |
| RMSD ($\text{\AA}$) | -0.32 | 0.399 | -0.33 | 0.380 |
| Hydropathy | -0.48 | 0.193 | -0.3 | 0.432 |

**Supplementary Table 8.**
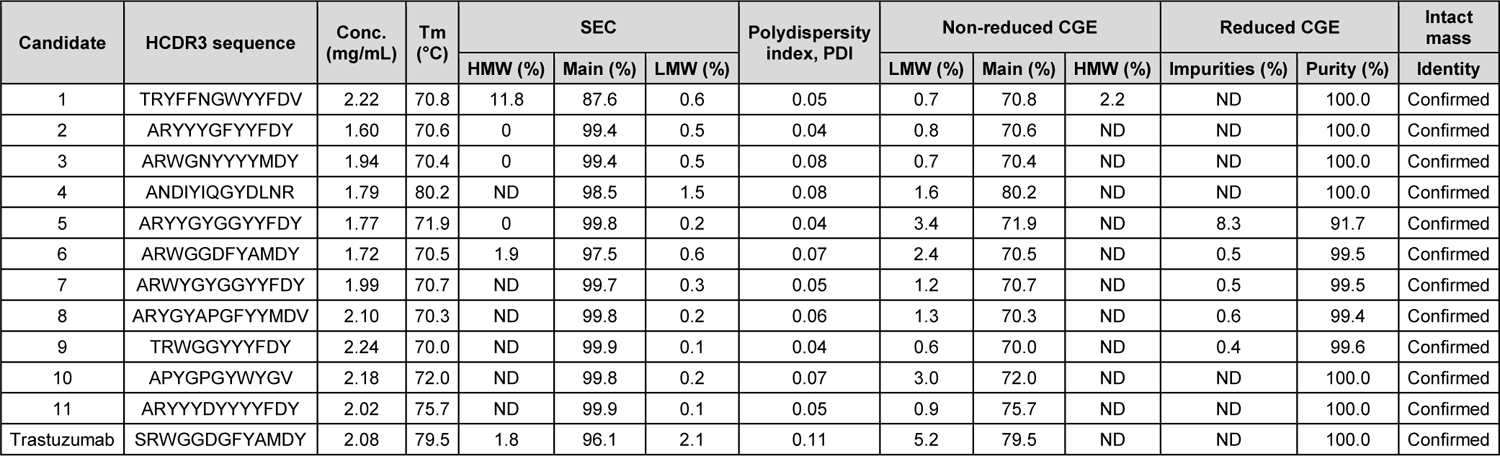
Product quality assessment of candidates reformatted as mAbs expressed in CHO cells. Determination of concentration, melting temperature (Tm), aggregates using size exclusion chromatography (SEC), colloidal particle size distribution (PDI), fragments using non-reduced capillary gel electrophoresis (non-reduced CGE), overall purity using reduced capillary gel electrophoresis (reduced CGE), and identity using deglycosylated intact mass for the 11 candidates reformatted into mAbs and trastuzumab produced by CHO cells. ND = not detected.

**Supplementary Table 9.**
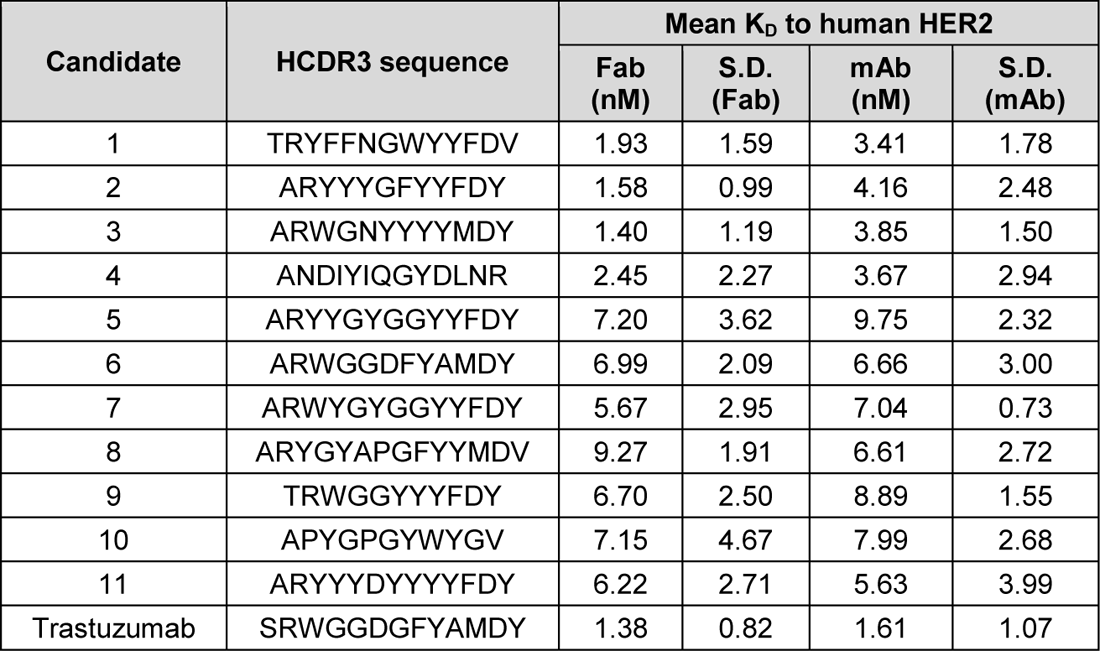
Comparison of the HER2 binding affinities of candidates as Fabs and as reformatted mAbs. SPR was used to determine the binding affinities of purified Fabs or reformatted mAbs of the 11 candidates and trastuzumab to human HER2. Experiments were run in replicates. The mean *K_D_* and standard deviations are presented. Correlations between Fab *K_D_* and mAb *K_D_* are Pearson R = 0.84 (*p* < 0.001) and Spearman Rho = 0.78 (*p* < 0.003).

**Supplementary Table 10.** Cross-reactivity kinetic parameters of candidates as reformatted mAbs. SPR was used to determine the binding affinities of the 11 candidate mAbs and trastuzumab to human HER2 as well as HER2 homologs from mouse, cynomolgus and canine. Binding affinities against family members HER1, HER3, and HER4 were also determined. Experiments were run in replicates. The single, high concentration injection of antigen (1 uM) and the mean *K_D_* in nM and standard deviations are represented. N/A = binding not detected.

| Candidate | Human HER2 | Mouse HER2 | Cynomolgus HER2 | Canine HER2 | Human HER1 | Human HER3 | Human HER4 |
| --- | --- | --- | --- | --- | --- | --- | --- |
| Trastuzumab | 6.2 ± 4.9 | N/A | 25.2 ± 12.5 | 7.8 ± 1.6 | N/A | N/A | N/A |
| 1 | 6.3 ± 4.1 | N/A | 49.8 ± 15.2 | 13.1 ± 3.0 | N/A | N/A | N/A |
| 2 | 4.6 ± 3.2 | N/A | 32.9 ± 6.8 | 7.6 ± 3.8 | N/A | N/A | N/A |
| 3 | 4.0 ± 4.3 | N/A | 4.6 ± 5.7 | 1.5 ± 1.1 | N/A | N/A | N/A |
| 4 | 4.8 ± 6.0 | N/A | 43.7 ± 34.5 | 4.8 ± 4.2 | N/A | N/A | N/A |
| 5 | 14.8 ± 3.3 | N/A | 254.2 ± 41.1 | 119.0 ± 6.7 | N/A | N/A | N/A |
| 6 | 8.9 ± 6.8 | N/A | 60.9 ± 23.8 | 18.5 ± 5.6 | N/A | N/A | N/A |
| 7 | 11.5 ± 4.9 | N/A | 171.0 ± 39.1 | 91.6 ± 18.7 | N/A | N/A | N/A |
| 8 | 9.0 ± 9.9 | N/A | 120.2 ± 9.8 | 37.6 ± 6.7 | N/A | N/A | N/A |
| 9 | 16.4 ± 5.9 | N/A | 198.7 ± 7.6 | 103.4 ± 7.9 | N/A | N/A | N/A |
| 10 | 17.5 ± 8.9 | N/A | 151.3 ± 20.9 | 5.0 ± 1.8 | N/A | N/A | N/A |
| 11 | 5.0 ± 4.4 | N/A | 104.1 ± 8.9 | 36.6 ± 2.4 | N/A | N/A | N/A |

**Supplementary Table 11.**
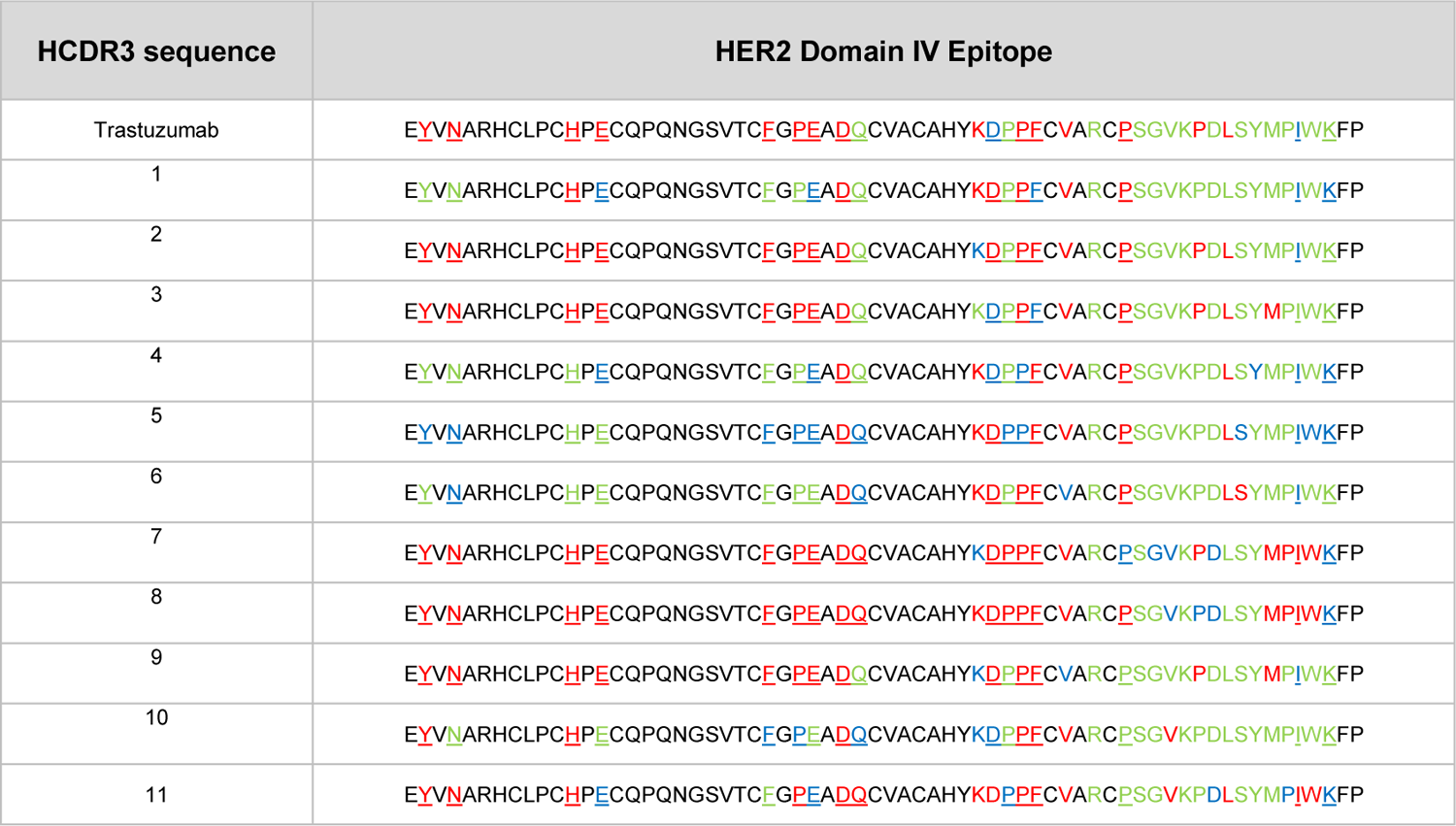
Domain IV epitope maps of the 11 candidate mAbs bound to HER2. Alanine scanning mutagenesis was used to mutate residues within 5 Å of trastuzumab Fab in PDB:1N8Z. Residues are colored according to their effects on HER2 binding by mAbs according to legend: green=not critical, blue=partially critical, red=critical. Residues that make important interactions with the heavy chain are underlined.

## Footnotes

1 We have open sourced the SPR binding affinity data as well as the functionality, developability, and cross-reactivity data from this study: https://github.com/AbSciBio/unlocking-de-novo-antibody-design.

a https://www.rosettacommons.org/software

b Motivation is given in the official Rosetta documentation for Fast Relax (https://www.rosettacommons.org/docs/latest/rosetta_basics/preparation/preparing-structures)

c https://github.com/AbSciBio/unlocking-de-novo-antibody-design

d https://docs.scipy.org/doc/scipy/reference/stats.html

